# Hardware-Efficient Compression of Neural Multi-Unit Activity

**DOI:** 10.1101/2022.03.25.485863

**Authors:** Oscar W. Savolainen, Zheng Zhang, Peilong Feng, Timothy G. Constandinou

**Affiliations:** Department of Electrical and Electronic Engineering, Imperial College London, South Kensington Campus, London SW7 2AZ, UK; UK Dementia Research Institute (UKDRI) Care Research & Technology (CR&T) Centre, based at Imperial College London and the University of Surrey, UK

## Abstract

Brain-machine interfaces (BMI) are tools for treating neurological disorders and motor-impairments. It is essential that the next generation of intracortical BMIs is wireless so as to remove percutaneous connections, i.e. wires, and the associated mechanical and infection risks. This is required for the effective translation of BMIs into clinical applications and is one of the remaining bottlenecks. However, due to cortical tissue thermal dissipation safety limits, the on-implant power consumption must be strictly limited. Therefore, both the neural signal processing and wireless communication power should be minimal, while the implants should provide signals that offer high behavioural decoding performance (BDP). The Multi-Unit Activity (MUA) signal is the most common signal in modern BMIs. However, with an ever-increasing channel count, the raw data bandwidth is becoming prohibitively high due to the associated communication power exceeding the safety limits. Data compression is therefore required. To meet this need, this work developed hardware-efficient static Huffman compression schemes for MUA data. Our final system reduced the bandwidth to 27 bps/channel, compared to the standard MUA rate of 1 kbps/channel. This compression is over an order of magnitude more than has been achieved before, while using only 0.96 uW/channel processing power and 246 logic cells. Our results were verified on 3 datasets and less than 1% loss in BDP was observed. As such, with the use of effective data compression, an order more of MUA channels can be fitted on-implant, enabling the next generation of high-performance wireless intracortical BMIs.

## 1 Introduction

### 1.1 Wireless Intracortical Brain-Machine Interfaces

BMIs are devices for connecting electronics to the nervous system. They function as a parallel nervous system, bypassing injuries in the nervous system or other obstacles, and are used in treatment of a wide range of conditions, from paraplegia, to paralysis and Locked-in syndrome [1], and more [2, 3]. BMIs come with different levels of invasiveness, where intracortical BMIs are the most invasive with electrodes being placed directly into brain tissue [4]. As a result, they give the highest spatial and temporal resolution of neural data, where they can measure the firing rates of individual neurons in the local vicinity of the electrode.

Historically, intracortical BMIs have used physical percutaneous connections, e.g. wires breaching the skin. These connect the electrodes in the brain to an external device that decodes the neural signals, typically a computer [1–3]. However, these percutaneous connections introduce significant infection and mechanical risks. They also degrade the quality of life of the user. This is why the move to wireless communication and powering is essential for the effective translation of intracortical BMIs into clinical applications, and it is one of the remaining bottlenecks [4–6]. In Wireless intracortical BMIs (WI-BMI), a small implant, placed on the brain or inside the skull, communicates wirelessly with a device outside the body, where further computationally intensive processing or decoding is done. Additionally, having millimeter or sub-mm scale free-floating WI-BMIs would be optimal in terms of minimizing potential injury and foreign body response in the brain [4].

### 1.2 Multi-Unit Activity

The most common signal in modern WI-BMIs is the Multi-Unit Activity (MUA) signal. When a neuron fires in the vicinity of an electrode, the electrode inductively measures a short, sharp voltage spike. MUA consists of measuring the timing of these spikes, assigning each spike on the same electrode to the same putative neuron, and binning the result at some temporal resolution, i.e, the binning period (BP). As such, each electrode channel outputs an integer value which is updated every BP, where the number represents how many spikes occurred on the channel in the BP. A typical BP is 1 ms, since spikes last approximately 2 ms. At a 1 ms BP, assuming only 1 bit per sample, that corresponds to a bit rate (BR) of 1 kbps/channel. This is a significant improvement over the raw broadband signal, which is typically sampled at some 16 bits/sample at 20 kHz, giving a BR of approximately 320 kbps/channel. The MUA extraction process, along with the extraction process of other common intracortical signals and their typical BRs, is given in Fig. 1 in the context of a typical WI-BMI data flow. The MUA signal is the most common signal since it is well-understood, easy to extract [7], has good decoding performance [8] and has relatively low bandwidth.

**Figure 1:**
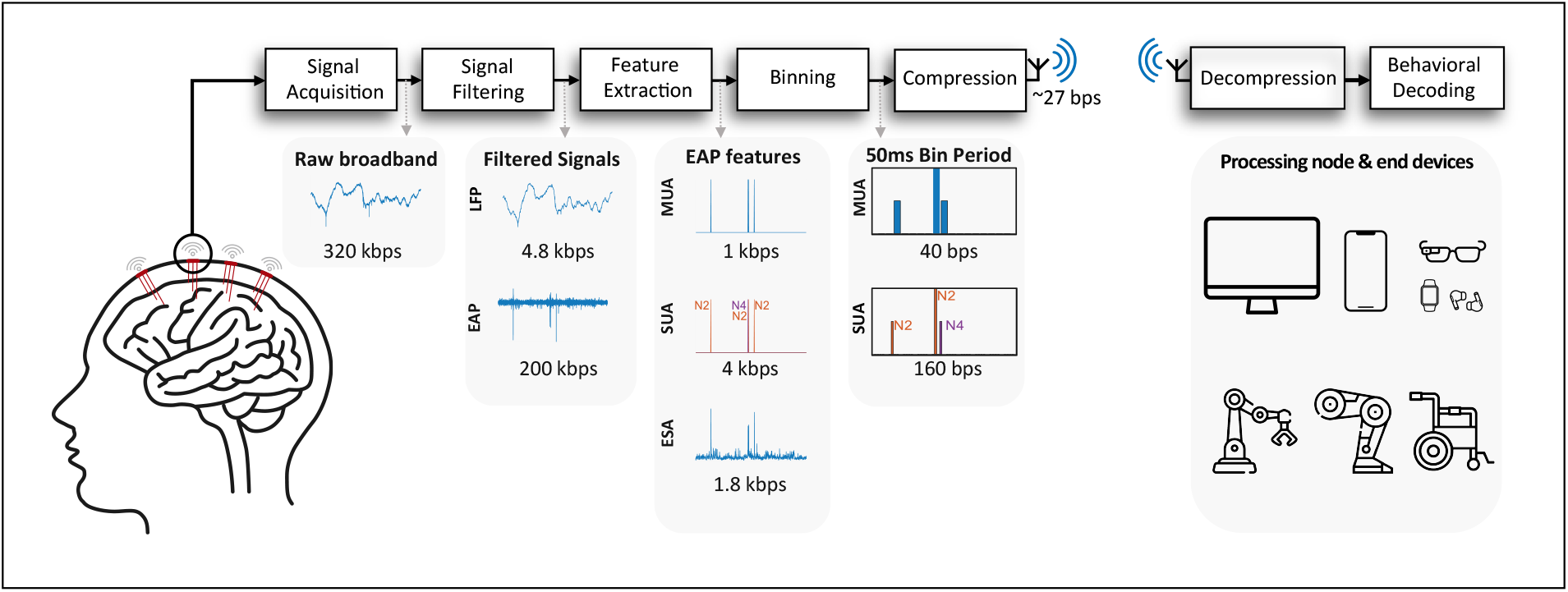
Typical BMI data processing and compression flow, with common extracted features / lossy compressions of intracortical broadband data. The numerical values beneath the signals give approximate BRs per channel for that signal. LFP: Local Field Potential, equal to the lowpassed broadband at approx. 300 Hz. EAP: Extracellular Action Potential, equal to the highpassed broadband at approx. 300 Hz. MUA: Multi-Unit Activity, equal to the unsorted thresholded spike activity from the EAP. SUA: Single-Unit Activity, equal to the sorted thresholded spike activity from the EAP. ESA: Entire Spiking Activity, equal to the enveloped rectified EAP signal, giving an envelope of unsorted spiking activity.

### 1.3 Heating in WI-BMIs

Heating is a major issue with WI-BMIs. It is known that heating tissue can cause irreparable damage [9]. This places important limits on acceptable heat dissipation into tissue from cortical implants, however safe levels of heat dissipation into cortical tissue are still poorly understood [9, 10]. The common standard is that power is strictly limited in WI-BMIs to an approximately 1 °C temperature increase or 1.6 mW/g of specific absorption rate (SAR) in tissue [4, 9, 11]. In the context of heating due to absorption of radio frequencies (RF), the IEEE standard C95.1-2019 gives limits for SAR heating depending on RF frequency [12]. However, it specifies that the understood SAR limits in brain tissue are generally derived from models and lack rigorous studies in live animals or humans, with significant variance between models [12]. FDA regulations further dictate that the local heat increase of brain tissue due to intracortical implants should be limited to only 0.5 °C [13]. In muscle and lung tissue, it is understood that up to 40 mW/cm^2^ heat flux can be allowed, however the limit is likely lower in cortical tissue [9]. We will assume a maximum heat flux limit of 10 mW/cm^2^ to hopefully provide a reasonable safety margin.

### 1.4 Communication Power

Communication power, alongside power management and the front-end ADC, is one of the most power-hunger modules for wireless implants. Therefore, there has been significant interest in lowering the communication energy per bit to keep the implant within heat flux limits. The most common data communication schemes for WI-BMIs are implemented using different shift keying (amplitude, phase, on-off) [14–18]. They are the most power-efficient solutions for implants; however, their BRs are below 20 Mbps. The ultra-wideband uplink proposed in [19] achieved 46 Mbps with 118.3 pJ/bit.

The implantable microsystems that are designed and fabricated based on full-custom application specific integrated (ASIC) are more power-efficient than microcontroller and field-programmable gate array (FPGA) based solutions. However, the design process of FPGAs is significantly less expensive and easier compared to that of ASICs. FPGAs also benefit from increased flexibility for programming, which better accommodates algorithmic changes. As a result, it is typical for researchers to use FPGAs to validate the ASIC’s performance before full development of the ASIC. Therefore the system in this work is implemented in a FPGA target. As such, this work assumes a system communication energy of 20 nJ/bit [20], which is state-of-the-art for FPGAs (Table 1). The average communication power per channel can then be calculated from the BR, given in (bps/channel):

**Table 1:**
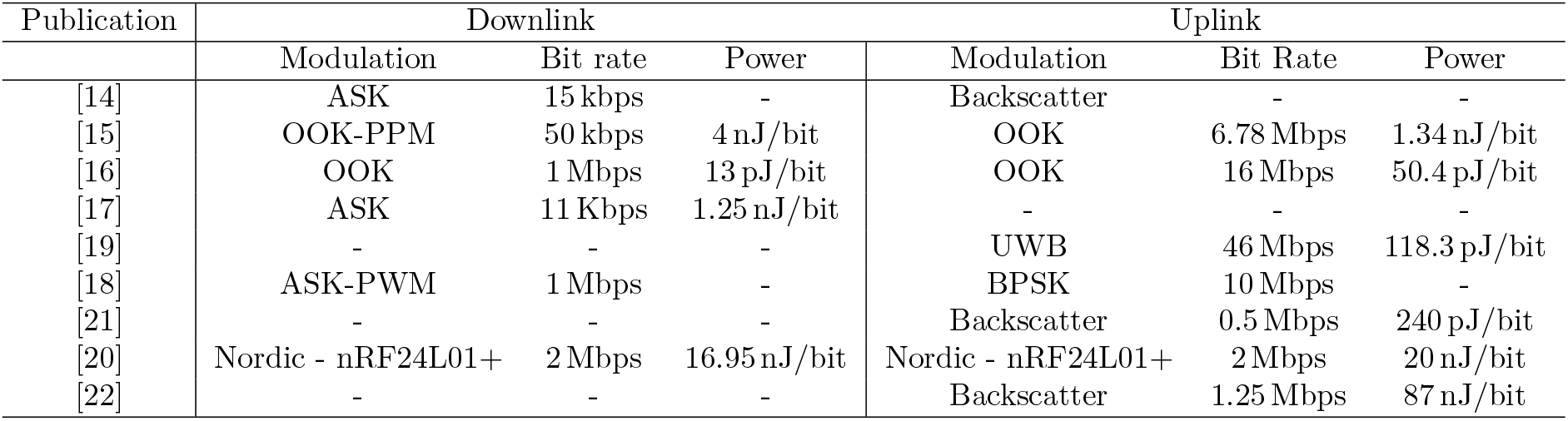
Wireless data communication power consumption comparison

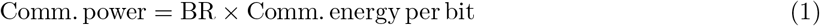

### 1.5 Data Compression in WI-BMIs

For a 10 mW/cm^2^ heat flux and a 2.5 mm × 2.5 mm scale implant, this gives a rough upper power budget of 625*µ*W. Assuming the entire implant power budget goes into communication and a communication energy of 20 nJ/bit for an FPGA-based WI-BMI [20], this translates into a maximum BR of 31.25 kbits/s. Therefore, such an implant could only communicate 31 MUA channels if all of the power was used for communication. This ignores that there are hardware static power requirements, as well as the front-end amplifiers, ADC, and any other necessary on-board processing to extract the MUA signal from the raw broadband recording.

Therefore, for the next generation of WI-BMIs, some form of data compression is necessary, even for MUA signals. Data compression comes in two forms. The first, is lossy compression, i.e. feature extraction. Moving from the raw broadband signal to MUA is a form of lossy compression, where information that is assumed to not be of interest to the final application is eliminated or degraded, saving on bandwidth. The second form of compression is lossless compression, where smaller codewords are given to more likely symbols; the basis of Information Theory [23]. However, many lossless compression algorithms consume significant hardware resources, e.g. Lempel-Ziv, Arithmetic, and Adaptive Huffman encoding.

In WI-BMIs, the available hardware resources are extremely few. As such, this work uses Static Huffman (SH) encoding, which is implemented in hardware as a small number of Look-Up Tables (LUT). The basic concept is that one pre-trains a SH encoder on representative MUA data and then implements it on the implant as unchanging LUTs that give Huffman codewords to the measured MUA data. If the SH encoder is appropriate to the data it encounters on-implant, it compresses the data. An example of a Huffman encoder is shown in Table 2. As long as the MUA data tends to have smaller firing rates (FR), the SH encoder in Table 2 will effectively compress the data. A SH encoder is as minimal an encoder as can be achieved in hardware, making it attractive for WI-BMIs. Additionally, Huffman encoding is optimal among algorithms that encode each symbol individually, making SH encoding effective while remaining simple. SH encoders were used in [24] to good effect to compress various intracortical neural signals (Entire Spiking Activity, Extracellular Action Potential, Local Field Potential) at different sample resolutions.

**Table 2:** Huffman encoder example. For the given frequencies and codeword lengths, the compressed data has an average length of 0.8 × 1 + 0.1 × 2 + 0.07 × 3 + 0.03 × 3 = 1.4 bits, versus the 2 bits that are normally required to represent 4 symbols with a fixed-length encoding.

| Huffman Encoder |  |  |
| --- | --- | --- |
| Symbol / Key | Frequency | Codeword |
| **0** | 0.8 | 0 |
| **1** | 0.1 | 10 |
| **2** | 0.07 | 110 |
| **3** | 0.03 | 111 |

### 1.6 Prior Work in the Compression of MUA

It was proposed in [6, 25] that increasing the MUA BP from the standard 1 ms may be an efficient way to lossily compress MUA data. This will be investigated in this paper. However, this compression is lossy because increasing the BP reduces the temporal resolution of the MUA firing times. This could cause two problems.

The first is increased delay in BMI-user experience. When we increase the BP, we increase the maximum possible lag between a neuron firing and the data being communicated off-implant. However, in [26], it was shown that the mean human reaction time in 120 healthy 18-20 medical students was larger than 220 ms for both auditory and visual stimuli. There are no other significant delays in the MUA WI-BMI data flow other than the spike counting, as the communication and other processing of the data occur on the ns scale. Therefore, a BP of 100 ms should be relatively well tolerated for motor decoding applications in terms of delay in user experience. From another perspective, it may also be worth considering that BMIs, with small BPs, may give users superhuman reaction times, which may be very desirable or even considered an unfair advantage in certain circumstances.

The second potential issue with increasing BP is reduction in Behavioral Decoding Performance (BDP). The reduced temporal resolution of neural firing times may reduce our decoding ability. However, the effect of varying MUA BP between 1 and 150 ms on BDP was investigated in [25]. The BDP was measured as the mean Pearson correlation coefficient between the actual and predicted X and Y axes of a free hand movement in non-human primates. It was found that increasing BP had a slight but statistically significant negative effect on BDP. The BDP reduced by 0.85% per 10 ms increase in BP, using a Long-Short Term Memory (LSTM) Neural Network (NN) decoder. Furthermore, [27] found that, for SUA signals, there was no difference in hand kinematic decoding ability between BPs of 10-100 ms for LSTM, Feedforward NN, and Wiener filter neural decoders. However, they also found that, when using Kalman filter decoders, increasing the SUA BP up to 50 ms improved the decoding, although not to the level of the NN decoders. Additionally, a 100 ms BP for motor decoding is a common choice by researchers [27–29].

As such, the effect of MUA BP on BDP is not clear, and it has been hypothesised that it likely varies by decoding algorithm and decoded task [6]. The effect of BP and limiting the dynamic range of MUA data on compression and BDP are one important aspect that will be further investigated in this paper.

### 1.7 This Work

In this work, we propose and compare multiple hardware efficient MUA compression schemes. As this is the first study on compressing MUA signals, we also evaluated how using one or multiple SH encoders, saturating the dynamic range, using on-implant histograms to add adaptivity into SH encoders, and setting different BPs can compress the data and affect the behavioral decoding quality. Our goal is to seek the best MUA compression algorithm with minimal resources to reduce data bandwidth so as to reduce the on-implant power (processing and communication power) without degrading BDP, while also keeping the signal temporal resolution within an acceptable range. The main contributions of this work are summarised below:

- Empirically showing the degree to which increasing MUA BP lossily decreases the communication bandwidth.
- Limiting the dynamic range of MUA data to reduce the communication bandwidth.
- The use of SH encoders for the compression of MUA data.
- The use of a sample histogram with mapping to add adaptivity to SH encoders.
- The use of multiple SH encoders, with assignment via a sample histogram, to add adaptivity to SH encoders.
- A novel ML algorithm for the offline selection of the best combination of SH encoders.
- A holistic analysis of the effects of MUA data compression on total implant power, BDP, hardware requirements, and temporal resolution of output data in a extremely low-power FPGA target.
- The use of statistical analysis to calculate the maximum amount of channels that could be hosted on-implant within power budget limits, given the variable-codeword lengths.

The rest of this paper is structured as follows. Section 2 detailed describes the dataset used in this work and different compression schemes. Section 3 shows the results of compression, decoding, and related hardware power consumption and resource usage. Trading-off among these metrics, a recommended setting is given. Section 4 discusses some design consideration based on the results and Section 5 concludes this paper.

## 2 Materials and Methods

The public datasets were loaded with Python 3.8 and MATLAB 2020a, the analysis was performed in Python 3.8, and the FPGA design in Modelsim Lattice Edition and iCEcude2 2020. The analysis Python code and FPGA Verilog code and designs have all been made publicly available at [30]. The formatted data and results have been made available at [31]. Researchers can use these to select their own compression system depending on their overall system requirements.

### 2.1 Datasets

To get a broad sample of MUA conditions, three publicly available datasets were used. These are summarised in Table 3, and further details are given in Supplemental Material, Section 2. For each dataset, the SUA data was intra-channel collated to MUA, then binned to the desired BP. The behavioral data was resampled to the same BP resolution using linear interpolation.

**Table 3:** Dataset summaries.

| Dataset | Neural data type | Species, electrode type and brain region | Details | Behaviour |
| --- | --- | --- | --- | --- |
| Flint [28] | SUA | Rhesus macaque monkey<br>Utah array<br>M1 | One subject<br>12 recordings across 5 days<br>96 channels<br>Recording lengths (quartiles, s):<br>597, 604, 630 | Free-reaching hand task<br>Continuous data stored |
| Sabes [32] | SUA | Rhesus macaque monkeys<br>Utah array<br>M1 and S1 | Subjects Indy and Loco<br>37 recordings for Indy across 10 months<br>10 recordings for Loco across a month<br>96-192 channels<br>Recording lengths (quartiles, s):<br>Indy: 472, 524, 816<br>Loco: 1771, 1928, 2384 | Free-reaching hand task<br>Continuous data stored |
| Brochier [33] | SUA | Rhesus macaque monkeys<br>Utah array<br>M1 and PMv/PMd | Subjects N and L<br>Single session recordings<br>96 channels<br>Recording lengths (s):<br>N: 1003<br>L: 709 | Hand reaching task<br>Target stored |

#### 2.1.1 Behavioral data

In this work, the X and Y-axis cursor velocities in the hand reaching tasks were used as the observed behavioral data. The BDP is defined in this work as the across-axes average Pearson correlation coefficient *r* between the predicted and observed X and Y-axis velocities. In the Brochier et al. dataset, the behavioural data consisted of labelled actions. As these were not continuous measurements, the behavioral decoding for the Brochier et al. dataset was not analysed in this work so as to keep the BDP metric consistent. The BDP metric is given in Eq. 2:

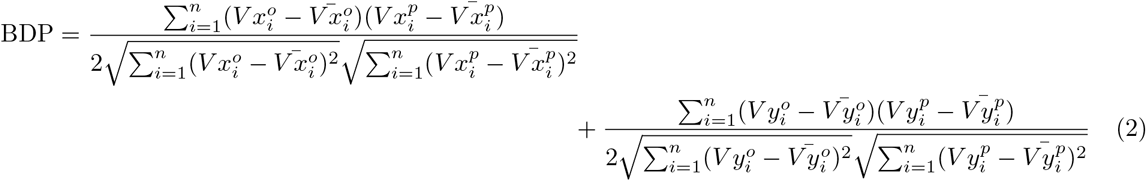

where 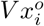 and 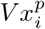 are the observed and decoder-predicted X-axis cursor velocities at sample *i*, 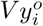 and 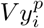 are the equivalents for the Y-axis, and *n* is the number of samples in the recording.

#### 2.1.2 Training-testing data split

The data was split into training (*A*) and testing (*B*) data. This was because many different systems with different parameters were considered, e.g. different BPs and *S* values, different module combinations, etc. Therefore, the training data was used to identify a well-performing system. The final system was then tested on the test data *B* so as to give an unbiased estimate of the system performance on new data.

The Flint dataset was split so that the first 4 days of recording sessions were included in *A*. This corresponded to 10 out 12 recording sessions. The remaining 2, taking place over another day, were used as testing data and included in set *B*. The Brochier dataset was all included in the testing data *B*. Finally, the Sabes dataset was split so that data from subject Indy was included in *A*, and the data from subject Loco was included in *B*. This was done so that the testing data included data from completely new subjects. This strengthened the test data, allowing us to test the system on new subjects to see if the BR performance and BDP were as desired.

### 2.2 Compression Modules

The full system overview is given in Fig. 3. Different module combinations (Detailed in Section. 2.2.5) were investigated and a full grid search of all system parameters was performed. We investigated each system in terms of BDP, temporal resolution, hardware resources and on-implant power consumption. That allowed us to analyse and trade-off among different metrics of interest for a WI-BMI so as to identify the best configuration. Finally, we tested the selected configuration on neural data from new subjects, and confirmed its compression and behavioral decoding performance. Details of the modules are given below.

**Figure 3:**
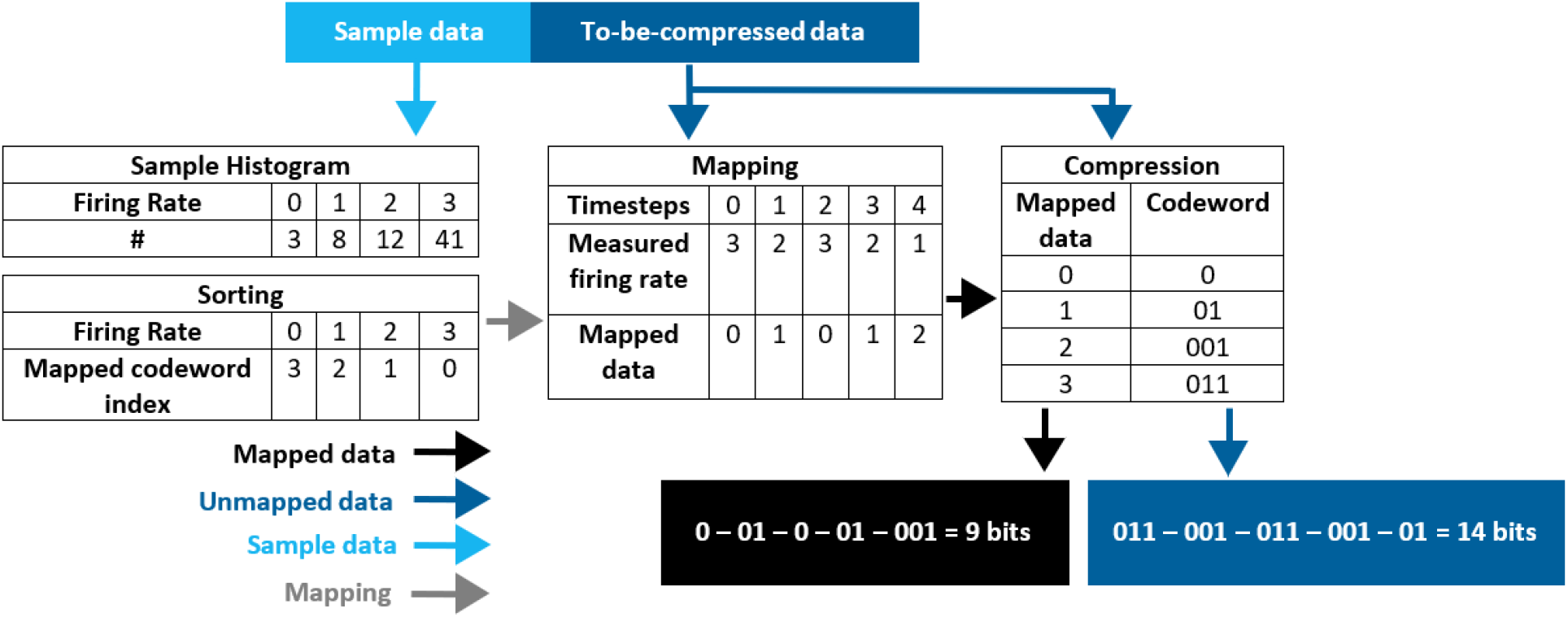
Use of a sample histogram to improve bit rates. A sample histogram is derived from the beginning of each channel’s recording. It is then sorted using a hardware-efficient sorting, where smaller indices are given to more common firing rates. The sorting is stored as a mapping, used to sort the rest of the data, i.e. the to-be-compressed data. The data is then compressed after mapping, where if a firing rate was found to be the x^th^ most common in the sample histogram, it was given the x^th^ shortest codeword. As such, the data histogram is approximated by taking a sample, and if the sample is well-representative of the rest of the data, this may help shorter codewords be given to more common firing rates, improving compression. In the example we can see the mapped compressed data requires only 9 bits, relative to the unmapped compressed data which requires 14 bits.

#### 2.2.1 Binning and saturation

Two lossy compression steps have been applied to compress the MUA data. Binning the MUA data at a certain BP to obtain the FR is a primary means of compressing the MUA data. We also investigated saturating the FR at a maximum value *S* − 1 to limit its dynamic range, where all FRs *>* (*S* − 1) were set to (*S* − 1). This means that there are fewer FR values that can be communicated, reducing the communication bandwidth. In order to test how different BP and *S* values can affect the compression and decoding performance, BPs of {1, 5, 10, 20, 50, 100} ms and S values of {3, 5, 7, 9} were tested.

These two operations perform a lossy compression to MUA signal, and therefore the degree to which they degrade the BDP was evaluated. The method is described in Section. 2.3.

#### 2.2.2 SH encoding

Applying SH encoders can losslessly compress the MUA data. As in Table 2, the idea is to give shorter codewords to more common values in an extremely hardware-efficient way. In this case, we give shorter codewords to more common FRs. The SH encoders were of length *S*, i.e. they had *S* input values and output codewords.

SH encoders need to be trained before use on representative data. Based on our observation on various recordings, we found that the firing rate distribution on average followed a decaying exponential, where smaller FRs were more common than larger FRs. This is shown in Fig. 2. As such, we trained the SH encoders on a decaying exponential, so they gave shorter codewords to smaller FRs. Further details are given in the Supplemental Material, Section 3. The SH encoder is represented by the ‘Encoder(s)’ block in Fig. 4. The use of multiple SH encoders will be discussed soon in Section 2.2.4.

**Figure 2:**
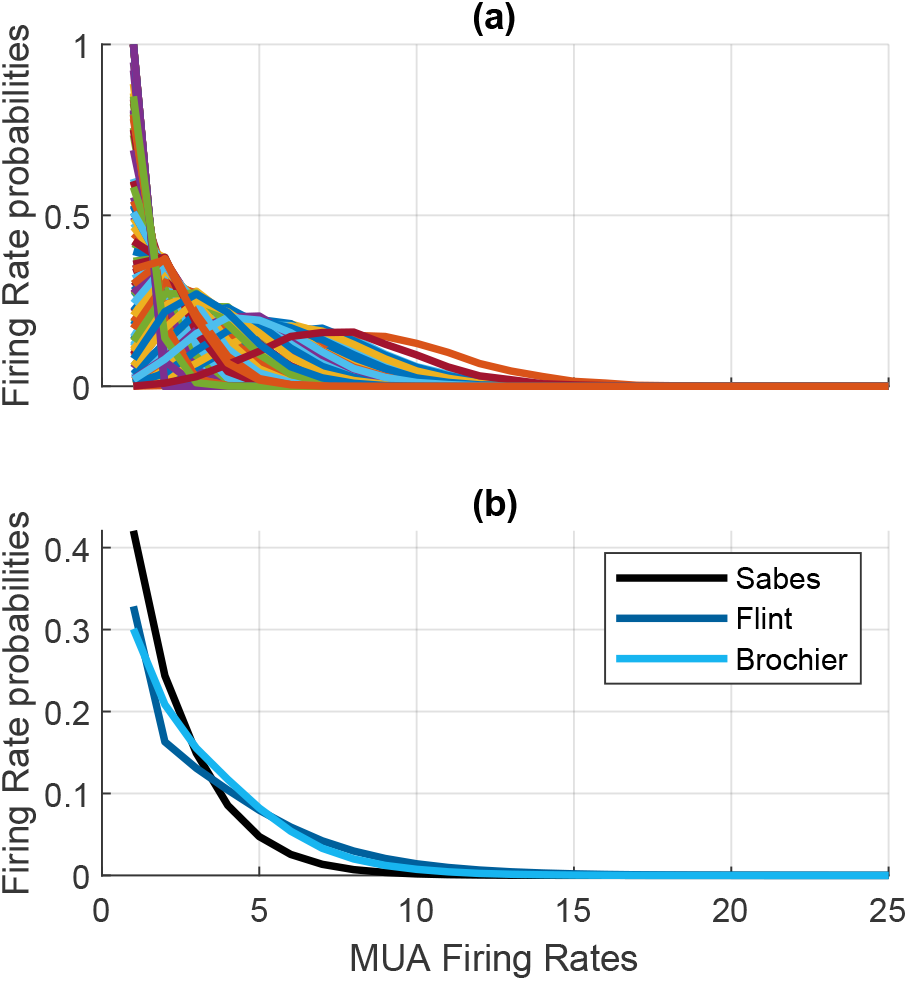
(a) A random sample of 100 channels’ MUA FR probability distributions with a 100 ms BP. (b) Average MUA FR probability distribution for each analysed dataset, with a 100 ms BP.

**Figure 4:**
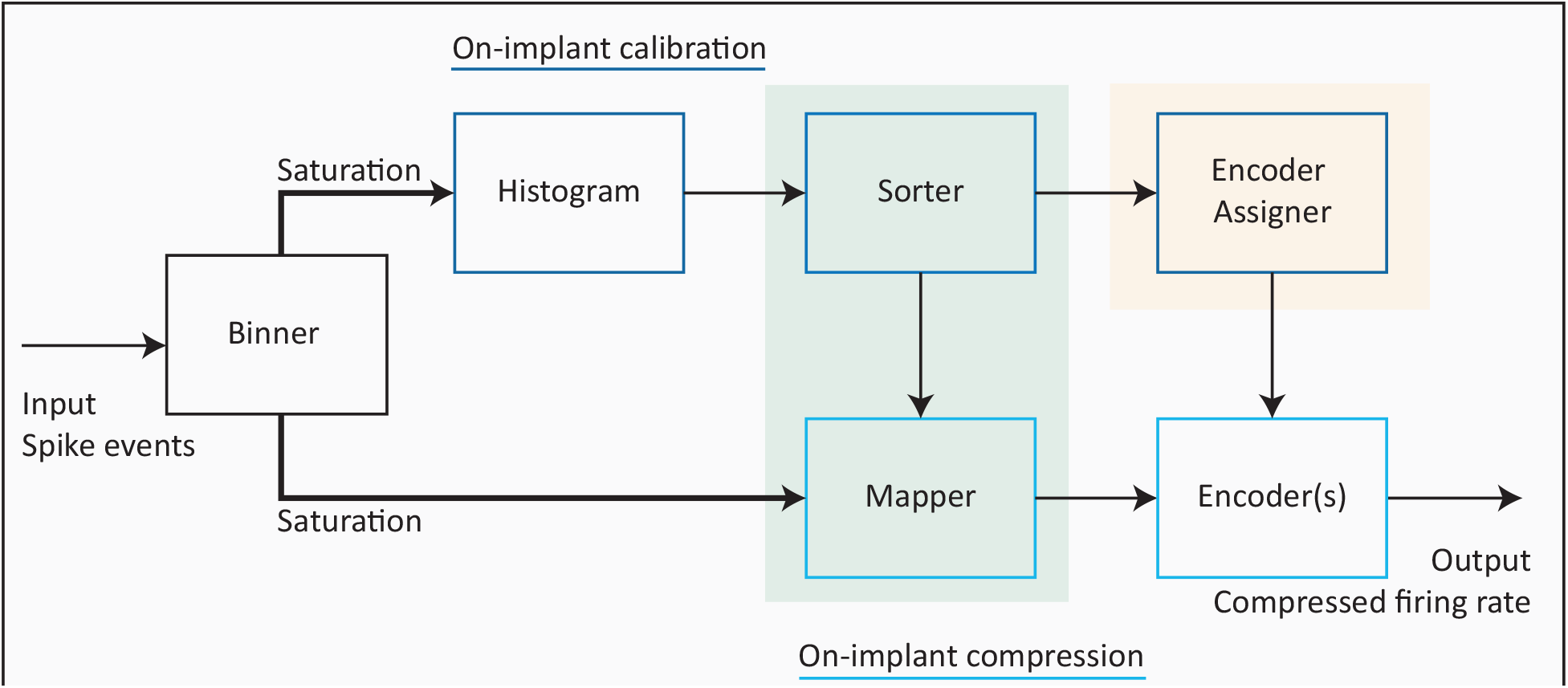
Compression data flow for each configuration. In the full system, a portion of the data is used for on-implant calibration, i.e. used to train a sample histogram for mapping and encoder selection. The mapping and selected encoders are then transferred to the main compression data flow, where the rest of the data is compressed. In the version without sorting and mapping, i.e. the ‘Without Mapping’ configuration, the green shaded modules are removed. If only *u* = 1 Huffman encoder is used on-implant, then the orange shaded module is removed, as no assignment is necessary. Finally, in the ‘Only Binning’ configuration, no on-implant calibration is performed, and the data goes straight from binner to transmission without a encoder.

#### 2.2.3 Firing rate mapping using histogram

As shown in Fig. 2 (b), for BP ≤ 100 ms, smaller MUA FRs are on average more common than larger ones. However, as can be observed in Fig. 2 (a), this is not always the case for each individual channel. As such, assigning shorter codewords to smaller FRs will not always give optimal compression. Here we investigate the use of a sample histogram to address this problem. The beginning of each channel’s recording was used to fill a sample histogram. This histogram was then used to estimate the relative frequencies of the FRs for each channel. The most common FRs in the histogram were then, for the rest of the data in each channel, assigned the shortest codewords via a hardware-efficient sorting (Supplemental Material Section 6.3). This was referred to as mapping the most common FRs to the shortest codewords, given the sample histogram estimate. As such, some semi-adaptability was introduced into the SH encoders. A demo histogram sorting and mapping process is represented in Fig. 3.

This module is represented by the ‘Histogram’, ‘Sorter’ and ‘Mapper’ blocks in Fig. 4. We considered histogram sizes of *d* = {0, 2, 4, 6} bits/bin, where there were *S* bins. Once 2^d^ samples had been measured, the histogram was considered to be full and was then used to estimate the FR frequencies. In the case of *d* = 0 bits, no histogram, sorting or mapping was used.

#### 2.2.4 Utilising multiple SH encoders

Multiple SH encoders can be used to increase the on-implant compression adaptiveness. This works by using the sample histogram to estimate which encoder would compress each channel the best. Each channel is then assigned its optimal encoder. Such assignment was obtained by taking the dot product of the histogram and the Codeword Length Vector (CLV) for each encoder. Dividing the dot product by BP and 2^*d*^ gives the BR. As such, we assigned each channel to the encoder that gave the channel histogram the smallest dot product (i.e. BR).

To give more information, the CLV is a vector of integers that represent the length of each of the SH codewords. For example a SH encoder of

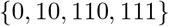

would have an CLV of

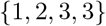

The dot product gives the total size of that channel’s communicated data after compression using an equivalent SH encoder.

For a SH encoder of size *S*, there are *h* possible non-redundant (i.e. with unique CLVs) SH encoders. We designed a custom Machine Learning (ML) offline algorithm to select the top co-performing *u* encoders from amongst all *h* possible encoders. In other words, this selected the best combination of *u* SH encoders by using offline MUA training data. This ensured that we had the best *u* encoders on-implant that channels could be assigned to, with *u ∈* [𝕫^+^, 1 ≤ *u* ≤ *h*]. The multiple on-implant SH encoders are represented by the ‘Encoder Assigner’ and ‘Encoder(s)’ block in Fig. 4. If *u* = 1 there was only one encoder on-implant, and so assignment was redundant since all channels went to the same encoder. We considered *u* values of {1, 2, 3, 5, 7, 10, 15, 20}.

The ML algorithm is further detailed in Section 3.3 of the Supplemental Material. It is an offline algorithm used in selecting which *u* SH encoders go on-implant, and is not itself present on-implant.

### 2.2.5 Module combinations

The modules included in each system configuration are given in Table 4. Similarly, the total parameter space investigated for the data compression is given in Table. 5.

**Table 4:** Required modules for each system configuration. The histogram is used for both assignment of encoders to channels, and for sorting/mapping of FRs. The system configurations vary if assignment is required or not, e.g. if only 1 encoder is considered, or if the histogram is to be sorted or not. The configuration also varies if no Huffman compression is considered, in which case only a binner and saturation are required.

|  | With Huffman encoding |  |  |  | Without Huffman encoding /<br>Only Binning |  |
| --- | --- | --- | --- | --- | --- | --- |
|  | With Mapping |  | Without Mapping |  |  |  |
| | $u > 1$ | $u = 1$ | $u > 1$ | $u = 1$ | | |
| Binner and Saturater | ✓ | ✓ | ✓ | ✓ | Binner and Saturater | ✓ |
| Histogram | ✓ | ✓ | ✓ | ✗ | Histogram | ✗ |
| Encoder Assignment | ✓ | ✗ | ✓ | ✗ | Encoder Assignment | ✗ |
| Encoder(s) | ✓ | ✓ | ✓ | ✓ | Encoder(s) | ✗ |
| Sorting and Mapping | ✓ | ✓ | ✗ | ✗ | Sorting and Mapping | ✗ |

**Table 5:**
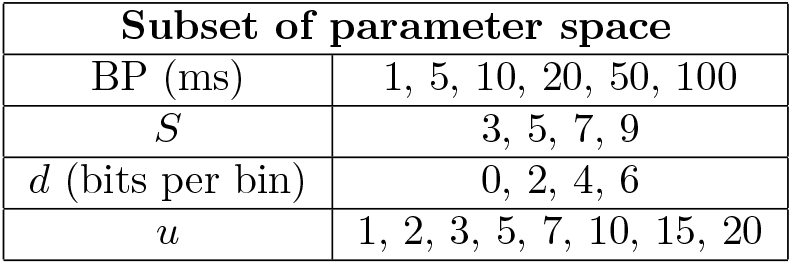
Considered subset of analysed parameter space.

| Subset of parameter space |  |
| --- | --- |
| BP (ms) | 1, 5, 10, 20, 50, 100 |
| $S$ | 3, 5, 7, 9 |
| $d$ (bits per bin) | 0, 2, 4, 6 |
| $u$ | 1, 2, 3, 5, 7, 10, 15, 20 |

There are in total 5 combinations. The first is the simplest system with only the binner and saturation, referred to as the ‘Only Binning’ system. The others use Huffman encoding, where the binned and saturated data goes straight to SH encoding. If multiple encoders are implemented, histogram and encoder assignment modules are needed in a calibration phase to select the best encoder for each channel. If we assume the FR data is not distributed according to a decaying exponential, the sorter and mapper are needed to map the more common FRs to shorter codewords. Therefore, the remaining four combinations are according to w/ or w/o mapping and *u* = 1 or *u >* 1. We refer to the one combination using all modules as the ‘Full System’. The FPGA implementation of the different modules is included in Supplemental Material, Section 6.

#### 2.2.6 Communication power estimation

For each parameter combination in Table. 5, we measured the compressed data BR using the training data*A*. From the BR, we derived the communication power from Eq. 1.

### 2.3 Determining the Impact of Lossy Compression on Behavioural Decoding Performance

Lossy compression involves losing information. In this case, the lossy aspects are increasing the BP, which reduces the temporal resolution of the neural data, and decreasing *S*, which saturates the data at an FR of *S* − 1. It is important to ensure that the lossy compression does not lose key information needed for the final application. In this case, the final application is the behavioral decoding of hand kinematics, which is a standard BMI behavioral measure.

As such, to ensure that not too much relevant information is lost, a behavioral decoder was implemented. It was used to decode the hand X and Y-axis velocities, using the BDP metric from Eq. 2. The input to the decoder was the binned and saturated neural data, for BP values of {1, 5, 10, 20, 50, 100} ms. To be exhaustive in our behavioral analysis, *S* values from 2 to 59 were investigated for their effect on BDP. However, due to the impossibility of automating all of the hardware optimisation, only *S* values of {3, 5, 7, 9} were investigated for the compression work.

A Wiener Cascaded Filter (WCF) was used for the decoder. WCFs have been found to have good decoding neural performance relative to other simple decoders, although they have generally found to not be as effective as deep learning methods [8, 27, 34]. However, their training times are significantly shorter [34]. As such, in this work they were used to investigate the relationship between *S*, BP and BDP. In this work, the WCF code from [8] was used. 5-fold cross-validation (hyper-)parameter optimisation was performed, and the details given in the Supplemental Material, Section 4. Once the parameters were optimised for each

*S* and BP, the BDP was calculated for each combination using separate testing data.

## 3 Results

### 3.1 The impact of lossy compression

#### 3.1.1 The impact of lossy compression on bit rate

It was proposed in [6, 25] that increasing BP would lossily compress MUA data, but neither evaluated how efficient that compression is. To the best of the author’s knowledge, the degree of the effect is empirically shown for the first time in this work. The BR required to communicate a channel of binned data is equal to *m/*BP (bps/channel), where *m* is the number of bits required to represent the unsaturated dynamic range (*S*^*′*^) of the FR. There is an approximately linear relationship between the lossless FR dynamic range and BP (Fig. 5 (a)). This produces a positive, approximately logarithmic effect on *m* from increasing BP (Fig. 5 (b). As such, merely increasing BP decreases the communication bandwidth (*m*/BP), relative to a lower BP (Fig. 5 (c)).

**Figure 5:**
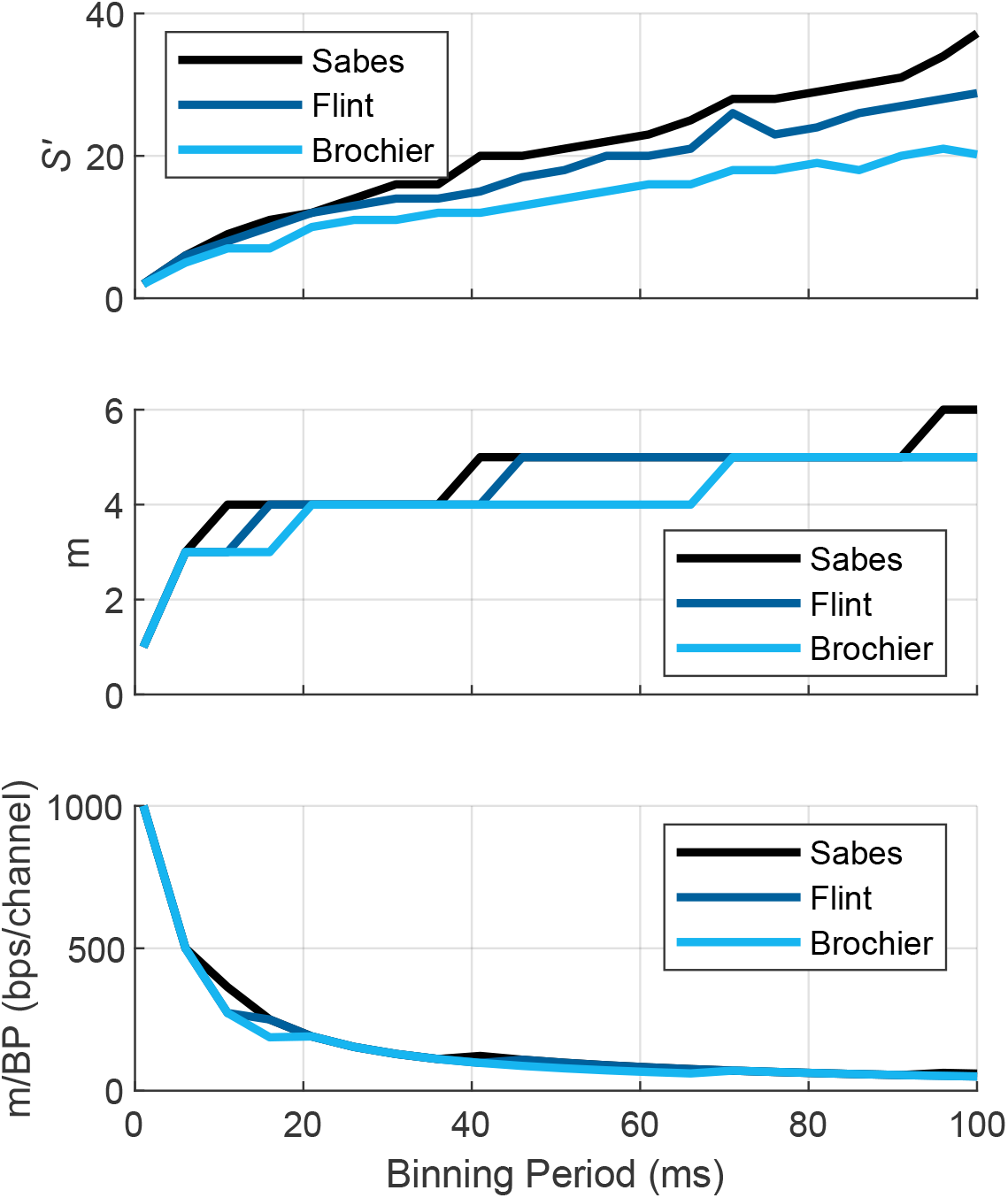
(a) A plot of *S*^*′*^ = *max*(*X*)+1, the dynamic range of the MUA FRs, as a function of BP, where *X* is the multi-channel MUA data. (b) The number of bits *m* = *ceiling*(*log*_2_(*S*^*′*^)) required to losslessly represent the dynamic range *S*^*′*^ as a function of BP, without any lossless compression. (c) The communication bitrate *m*/BP (bps/channel) required to communicate a dynamic range of *S*^*′*^. (a-c) The analysed MUA data *X*, from which *S*^*′*^ is measured, is the entirety of the training data *A*.

Saturating the FR range by setting the dynamic range *S* to *S < S*^*′*^ can obviously further reduce the BR, as the BR is proportional to the dynamic range to be transmitted. That is not the only advantage of saturating. If a SH encoder is used, a large glossary of the possible FRs to be compressed means a large SH codebook. Limiting the max FR from 10s to less than 10 can significantly reduce the size of the on-implant SH encoder and therefore reduces the power and resources.

#### 3.1.2 Impact of lossy compression on BDP

Fig. 6 (a) shows the BDP vs. BP and *S* results averaged across the Flint and Sabes datasets. Examples of observed and predicted behavioral data are shown in Fig. 6 (b-d).

**Figure 6:**
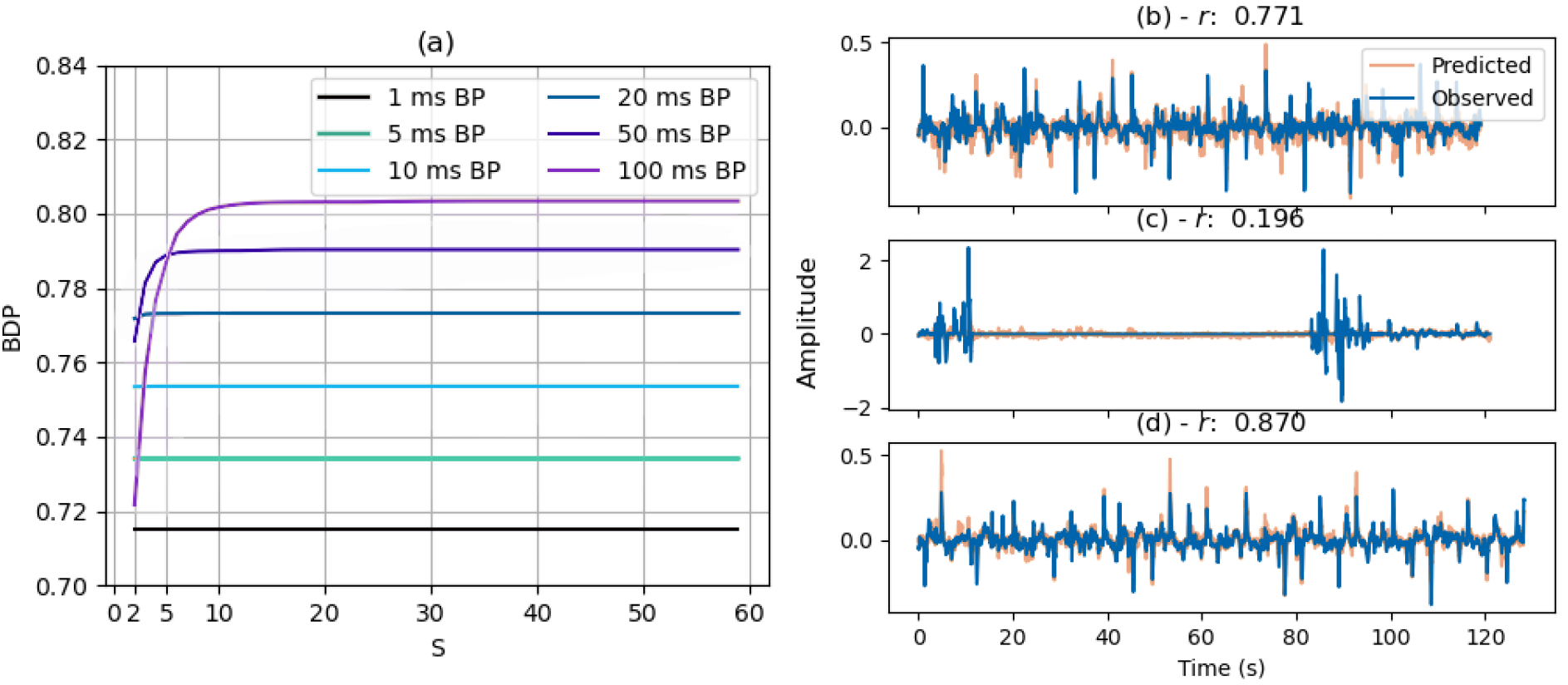
(a) Behavioral decoding performance (BDP) as a function of BP and *S*. Each S/BP combination was parameter optimised on 5-fold CV, with the results averaged from the Sabes lab and Flint datasets. (b-d) Example observed vs. predicted X-axis velocities from 5-fold CV, with corresponding BDP (*r*) for random Flint recording and parameter combinations during parameter optimisation at a BP of 5 ms.

Fig. 6 (a) shows that the BDP improves as a function of BP, and is unaffected by *S* if *S* is large enough. For BP ≤20 ms, even *S* = 2, i.e. binary representation, is not lossy enough to affect the BDP. For BP at 50 or 100 ms, *S* = 3 or 5 are large enough to have less than 2% BDP degradation. For example, a 100 ms BP and an *S* value of 5 would give a BR of *ceiling*(*log*_2_(*S*^*′*^))/BP = 30 bps/channel, while using no lossless compression. A typical bit rate of MUA signal is 1 kbps. Therefore, a 33 times bandwidth reduction can be easily achieved with just the use of binning and saturation lossy compression, with minor to no degradation on BDP.

### 3.2 The impact of static Huffman encoding

The SH encoder is a lossless compression operation applied after binning and saturating. Fig.7 shows the reducing effect on BR from the addition of just a single SH encoder.

Overall, roughly another 50% bandwidth reduction can be achieved by using a single SH encoder. Moreover, as benefited from the saturation, the SH encoder with a small codebook can be implemented with only minor resources (less than 100 logic cells) consuming negligible power compared to the binner.

### 3.3 The impact of improving adaptiveness

Using an on-implant histogram to map FRs and select a suitable encoder from multiple on-implant encoders can improve the adaptiveness of the compression. This can be especially effective when the distribution of FRs to be compressed is unlike in the training data.

According to our results, using histogram to map FR can reduce the BR by another 5% to 20% as the histogram size increases. This improvement is only noticeable when the BP is 50 ms or 100 ms because longer BPs cause the FR distribution to vary more from the standard decaying exponential. In which case, increasing the histogram size estimates the true distribution more accurately, and so provides a more accurate mapping, making the effect of the histogram notable. However, the FR distribution of short BPs varies less, which makes the mapping mostly redundant. Mapping the FR with local information can even degrade the compression performance, regardless of BP, when the histogram size is too small. This is because if the beginning of the recording is not representative of the rest of it, our mapping may be maladaptive and perform worse than the assumed decaying exponential. As such, larger histogram sizes are more reliable.

Using more encoders does not bring significant improvement on BR. We noticed that the same encoder is nearly always selected even when there are multiple available encoders. This ‘best’ encoder was the one trained on a sharp decaying exponential, where the CLV was of form [1, 2, …, *S* − 1, *S* − 1]. That suggests, on the one hand, that our ML-based encoder training algorithm works well in selecting the best encoder; on the other hand, the FR distribution when BP is less than 100 ms does not vary enough to require the adaptiveness provided by having more than one encoder.

The adaptiveness however comes with hardware costs. The on-implant histogram, dot product and sorting are all resource-hungry. We have done intensive hardware optimisation and the details are provided in Supplementary Section 6 and 7, but their resource occupation can still be the bottleneck of the whole system.

### 3.4 System configuration selection

In previous sections, we have shown how different modules can contribute to the MUA data compression, and their cost on BDP and/or hardware. Next we will determine the best system configuration.

We finalised a configuration by empirically testing all combinations of different modules, BP, *S*, histogram size and the number of encoders shown in Tables. 4 and 5. We then traded off between the system behaviour decoding performance (BDP), behavioural decoding temporal resolution (BDTP, i.e. the temporal resolution we had of the neural data), hardware total power (communication + processing) and resource usage as shown in Fig. 8 (a). Each point stands for one parameter setting and different colors represent the system operating at different BP. The points high up on the resources axis indicate systems with larger *S*, histogram size or more on-implant encoders. The points that are perpendicular to the rest, with low resources but high power, are the setting with only the binner at different *S* values, as these had low hardware usage but higher communication power since no lossless compression was used.

**Figure 7:**
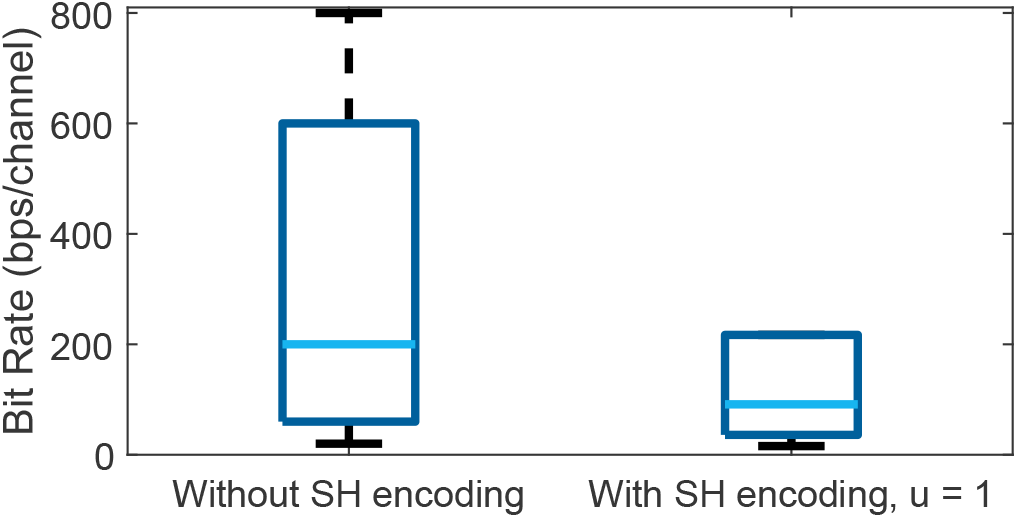
Boxplot of bit rates of all compression systems with and without Huffman encoding at *u*=1. This shows that the addition of a single SH encoder improves compression performance by more than 2 times on average.

**Figure 8:**
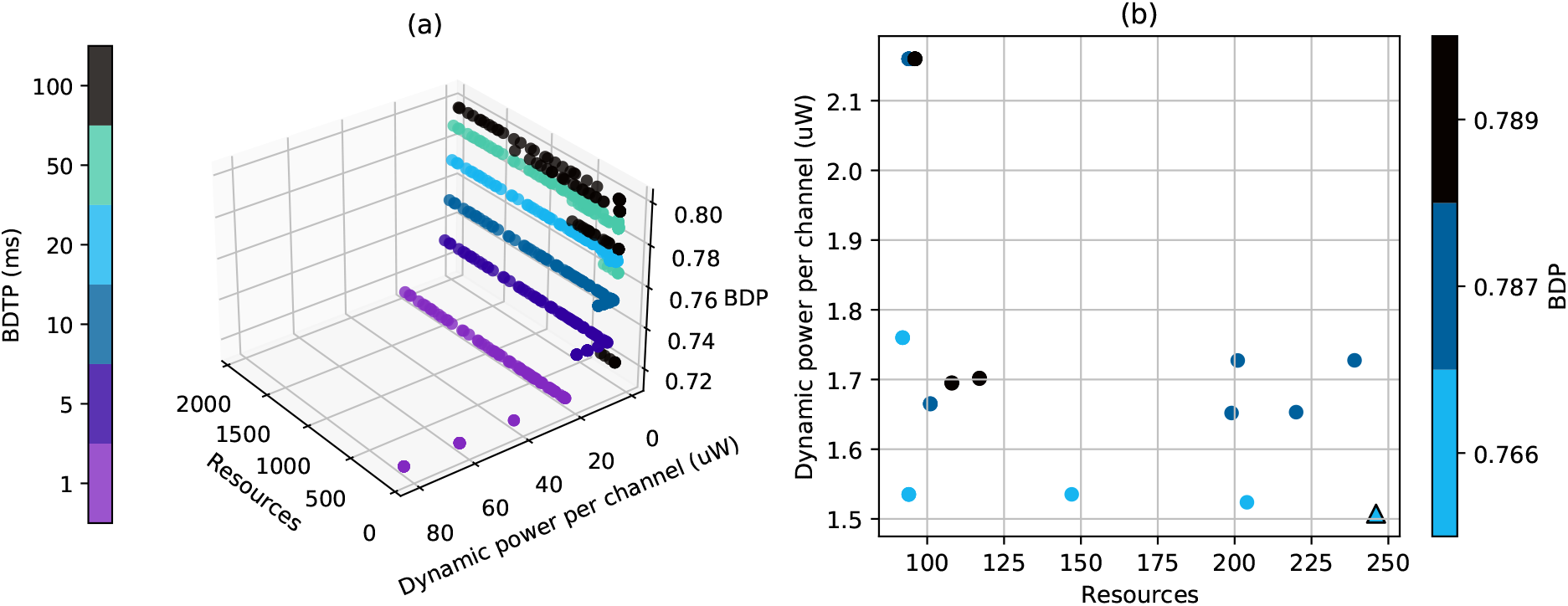
(a) Integrated results: BDP and BDTP values for different resources and dynamic power consumption levels. (b) Sample of integrated results, with BP/BDTP = 50 ms, resources *<* 260, dynamic power *<* 2.2 *µ*W/chan for our 128 channel system. The outlined triangle represents the chosen system configuration for our tested system. Note that the color bars for (a) and (b) are distinct.

The best trade-off is shown in Table 6. As this is the first system on MUA data compression and a massive parameter space is involved, it is worthy to provide some findings and considerations on how the system is selected.

**Table 6:**
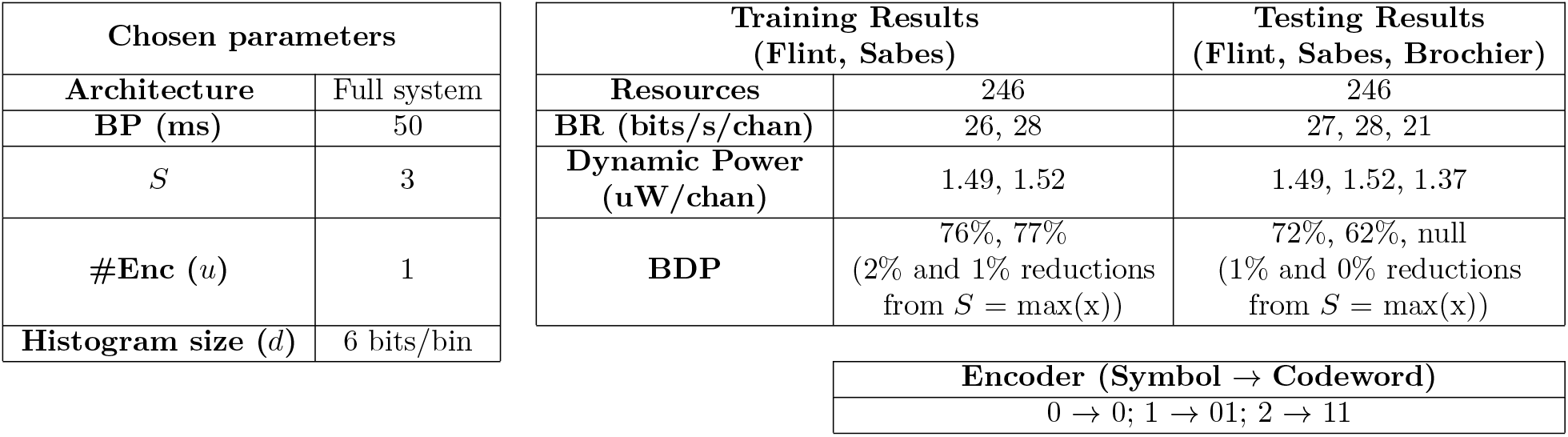
Chosen system parameters and encoder for testing and associated training (*A*) and testing (*B*) results. For the training and testing results, all data in *A* and *B* were respectively used.

#### 3.4.1 The selection of BP and S

BP in our design is directly equal to the temporal resolution BDTP. Increased BP or BDTP can bring higher BDP and lower communication power as the firing rates are transmitted less frequently. However, *S* needs to be sufficient for high BPs to enable sufficient BDP. It was observed that at a BP of 100 ms and *S* = 3 that the BDP suffered significantly. This can be observed in both Fig. 6 (a) and Fig. 8 (a) as a cluster of dark points (100 ms BP) with low BDP. As such, given sufficient *S*, increasing the BP is a very attractive prospect. It needs to be balanced with the desired temporal resolution of the decoded output, but BP should be able to be increased significantly without negatively affecting user experience, as discussed earlier in Section 1.6.

In our case of behavioural decoding, 50 ms can be a good choice. It is conservative in terms of the impact on user delay, while also having high BDP and low communication power. As such, we selected a BP of 50 ms. Furthermore, according to Fig. 6, when the BP is 50ms, limiting the FR dynamic range to 3 produces a 2% BDP degradation. As such we selected BP = 50 ms, *S* = 3.

#### 3.4.2 The selection of the number of encoders and histogram size

As previously stated, using more than one encoder does not significantly reduce BR and therefore is not considered. Using no SH encoder can be especially resource-saving. However, the approx. 50% bandwidth reduction brought by a single SH encoder can significantly reduce the communication power and therefore the total power. The points that belong to a perpendicular segment relative to the rest of the data in Fig. 8 show how the addition of SH can significantly reduce communication power.

With respect to the histogram size, though it needs more resources, it improves the compression while the resource usage is still acceptable. To get more insight, we focused in on Fig. 8 (a) with BDTP = 50 ms, resources ¡ 260 and dynamic power ¡ 2.2 *µW* /channel, shown in Fig. 8 (b). The bottom four points from left to right are the configurations with one encoder, *S* = 3, and histogram sizes of [0, 2, 4, 6] bits/bin respectively. It makes sense, in terms of scaling with channel count, to prioritise the lowest power configuration. This is because resources are roughly static at 246 with increased channel count, which is acceptable. Additionally, BDP increases somewhat logarithmically with channel count according to neuron dropping curves [35, 36], and BDTP is unaffected. However, power increases roughly linearly with channel count, making it the parameter that scales least well. Therefore as long as the resource usage is acceptable, reducing the power consumption should be the first priority. However, for resource-constrained scenarios, the bottom left setting can be selected which is the configuration without histogram, i.e. no mapping, where the encoder compresses the binned firing rate directly (for data flows see Fig. 4).

### 3.5 Testing data results

Next, we looked at testing our chosen system on data it had not seen yet. Given our chosen system parameters, summarised in Table 6, we determined the BR and BDP on the testing data *B*, also shown in Table 6.

#### 3.5 1 Communication power results

For the Flint, Sabes and Brochier data in *B*, the across-channel-and-recording average BRs were 26.5, 27.8, and 20.6 bps/channel respectively, corresponding to dynamic power/chan values of 1.49, 1.52 and 1.37 *µ*W. These are highly similar to those in the training data, where the averages for the Flint and Sabes data were 26.4 and 27.8 bits/s/chan and 1.49 and 1.52 *µ*W respectively. This is especially significant for the Sabes and Brochier test data, which consisted of subjects that were completely separate from the data in *A*. This means that the system effectively compressed data from 3 entirely new non-human primate subjects. This can be compared to the 40 bps/channel produced by the Only Binning configuration where no Huffman encoder is used (ceiling(log_2_(3))/(50 ×10^*−*3^)).

#### 3.5.2 Behavioral decoding results

The BDP was measured for the Sabes and Flint datasets using the testing data in *B*. Each channel was split 90-10% into training and testing sets. *S* and the BP were fixed at 3 at 50 ms respectively, and as in Section 2.3 the WCF hyper-parameters and pre-processing parameters were 5-fold cross-validated on the training set. The best parameters for each BP/*S* combination were taken, and the BDP measured on the testing set.

The average BDP for the Flint dataset was 0.724, and for the Sabes data it was 0.616. These BDP values from *B* are significantly lower than in the training data *A*. At first glance this is worrying, since it may suggest the compression scheme was overly lossy and too much behavioural information was lost. However, close examination of the results indicate that the tested system’s BDP values are lower because the recordings have less behavioral information in them, or the recording quality is less good, etc. This is shown by comparison of Fig. 5 and 8 in the Supplemental Material, as well as Fig. 6 in the main manuscript. In particular for the Sabes test data, they show that the data seemed to suffer from a few recordings with very low BDPs (e.g. at approx. 0.4), which dragged down the average to significantly below the median BDP. A larger amount of Sabes test recordings performed quite well (approx. 0.7), although they still generally performed worse than the train *A* data. As such, the test recordings seem to simply be of worse quality than the train recordings.

Furthermore, across all test recordings, the tested system’s compression did not negatively affect the BDP by more than 1.62% compared to the top observed BDP for each recording across all BP and *S*, and for the overwhelming majority of recordings the impact was 0. This is encouraging, as it shows that the quality of the recordings was the decisive factor in the BDP, not the lossy compression.

The main takeaway for the BDP results is that reducing *S* to 3 for a BP of 50 ms had no significant negative effect on BDP. It is expected that if the recording quality is similar to that in the training data, higher BDPs will result. It is also likely that using more advanced deep learning decoders would result in higher BDPs [8, 25, 27].

## 4 Discussion

### 4.1 The number of channels supported by the power budget

As discussed in Section 1.5, for a 2.5 mm × 2.5 mm scale FPGA implant with a 625 *µ*W power budget, one could wirelessly transmit up to 31 channels if all the on-implant power was used for communicaiton. In practice, with a static FPGA power of 162 *µ*W, negligible spike detection power [7] and a processing power for the 1 ms binner of 0.96 *µ*W/channel, a maximum of 22 channels could be measured on-implant.

However, our chosen system’s power consumption depends on variable-length codewords. Therefore, there is a risk that the BRs will be higher than the expected ∼ 27 bps/channel as given in Table 6. For example, one might measure a handful of particularly active channels, and this may increase the BR. If one chooses the number of channels on-implant so as to be close to the permitted power budget, and the channels are more active than expected, this may produce more heat than desired. As such, it warrants choosing the number of channels based on a statistical understanding of the worst case scenarios.

As such, we took random samples of the channels. For sample size *z* ∈ ℤ^+^, we took 100,000 samplings of *z* random channels. For each random sampling *Y*, from the channels’ summed BRs we obtained the resulting total system power *P* using the estimates in Equation 3:

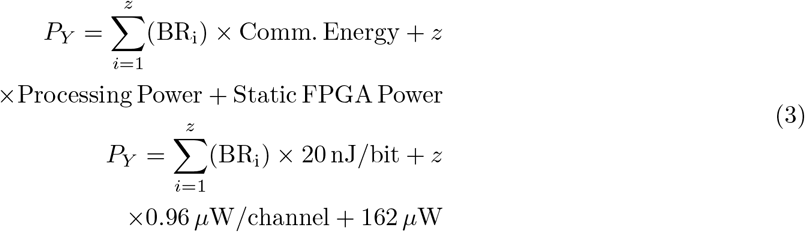

where BR_i_ is the BR of the i^th^ channel in sampling *Y*, where 1 ≤ *i* ≤ *z, i* ∈ ℤ^+^.

It was then determined, for each number of channels *z*, what percentage of random channel combinations exceeded the desired power budget *B*:

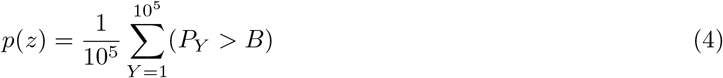

where (*P*_*Y*_ *> B*) is a boolean value equal to 1 if *P*_*Y*_ *> B* and 0 otherwise. As such, *p*(*z*) gave a permutation derived *p*-value for each number of channels *z* not exceeding the power budget.

Using our chosen architecture and power budget of *B* = 625*µ*W and averaged across our 30 CV runs, it was found that from the training data results that having up to 304 channels never exceeded the power budget. Having 305 channels had a *p*-value of ∼5e-4 of not exceeding the power budget with *p*(305) = 55*/*10^5^, and 306 channels or higher had significant chances of exceeding the power budget of *p*(*z >* 305) *>* 0.05. As such, we think having approximately 300 channels for our 2.5 mm ×2.5 mm FPGA hardware is ideal assuming the given power estimates hold true, while staying within a conservative heating safety margin. As such, by compressing the MUA data one can send out over 13 times as many channels as when sending out the raw MUA data for a similarly sized FPGA device. In ASIC, this difference would likely be far more pronounced, given the reductions in dynamic and static power. However, the contribution of the front-end amplifier and ADC would need to be included as they would likely be integrated. Given that ADCs with power consumption as low as 0.87*µ*W/channel have been achieved [37], there is reason to believe that impressive channel counts could be obtained at mm-scale in ASIC.

#### 4.2 Configuration selection considerations

The configuration selection can be application-dependent. It warrants mentioning that 100 ms BPs for behavioral decoding are common in the literature [6, 28, 29]. Therefore, a *>* 50 ms BP system could also be of interest. Additionally, if increased BDP is an absolute priority, then a configuration with a higher *S* may be appropriate. However, one should consider that if increased power is required as a result, this can reduce the amount of allowable channels for an implant of the same size, perhaps reducing the final BDP. Similarly, if a system has only a few recording channels, where communication power does not dominate, one may choose a system that significantly reduces hardware requirements over marginally reducing BR, e.g. a ‘Without Mapping’ configuration.

All results and hardware designs are made publicly available, and researchers are free to select from them for their own system designs.

### 4.3 The effect of BP on BDP

We found that BDP increases as a function of BP between 1 and 100 ms. Although this differs from some other results that used different decoders [25], it has also been theorised in the literature that we should expect BDP results to vary by decoded behavior and decoding algorithm [6]. [25] found that increasing BP reduced BDP, but it used Long-Short Term Memory (LSTM) neural network decoders, which are a form of deep decoder. It is unsurprising that a deep decoder that can exploit long-term temporal dependencies to find extra information in high-precision timing of neural firing rates compared to a simple WCF decoder. As such, the difference between the relationship between BP and BDP in Fig. 6 and [25] is not surprising.

### 4.4 Generalising the Results to Other Behaviors

How well lossy compression has performed depends on the final use of the data, and whether any key information has been lost. In this work, the final outcome was the decoded hand kinematics, which are a standard BMI behavioral measure [8,25,28,32,33,38]. Therefore, although the lossy aspect of the compression system is tailored to a specific task, it is a general task that is ubiquitous across BMI research. Additionally, the lossy aspect of the data compression scheme is very simple: increasing the BP, which is standard during BMI behavioral decoding [6, 28, 29, 38, 39], and saturating the MUA data, which had only a negligible effect on BDP for this task.

For this hand kinematic task, the system’s BDP was tested on a completely new subject, ‘Loco’ of the Sabes dataset, and for all tested recordings the BDP was at most negligibly reduced by data compression (1.6% in the worst case, 0% in most cases). For the Flint test recordings (new recordings on the same Flint subject as in the training data), the BDP was similarly unaffected by lossy compression. As such, we can say that the performance of the compression system was robust across different subjects, which is a very significant result. However, we cannot say the same across tasks. It is our belief that the results will probably be consistent across tasks decoded from the motor cortex, but if not, then *S* can simply be increased or BP varied.

Future work will look at decoding hand-writing kinematics, using the publicly available data from [2]. As such, so far we can only say that the tested compression scheme generalised very well to 3/3 new subjects in terms of compression, and to 1/1 new subject in terms of behavioral decoding for hand kinematic tasks in WI-BMIs.

### 4.5 Fixed Length vs. Variable Length Codewords and Bit-Flip Errors

It warrants mentioning that lossless compression works by giving variable length codewords to symbols. Due to the multiplexed encoding of MUA, this makes losslessly compressed MUA data more vulnerable to bit flip errors making the multiplexed communicated data block undecodable. As such, some noisy channel encoding or decreasing the BP may be necessary, assuming the bit flip error rate is sufficient to warrant it. This would increase the BR marginally, and is discussed further in the Supplemental Material, Section 8.

### 4.6 Multi-Channel Compression

Future work will consider compressing the entire MUA signal across channels, whereas this work looked at compressing intra-channel MUA. It may be that some dimensionality reduction is possible, or that correlations between adjacent channels can be taken advantage of as in [40] to further compress the data without reducing BDP or other metric of interest.

On-going work is looking at methods to compress the MUA in ‘asynchronous’ architectures, where the MUA FR is only communicated for a channel if it is larger than 0 for the given time period. Preliminary results show that, for BPs higher than or equal to 20 ms, the methods in this paper are superior. For BPs lower than 20 ms, asynchronous methods seems to perform best.

## 5 Conclusion

In conclusion, our objective was to reduce the MUA data bandwidth so as to reduce the MUA-based WI-BMI power consumption and prevent tissue heating and damage. We eventually achieved nearly 40 times MUA bandwidth reduction from 1kbps/channel to 27bps/channel with 2% decoding degradation on training data, and less than 1% on testing data. Such a distinguishable achievement is made by a binner at 50 ms BP, a dynamic range limited to 3 possible values, hardware efficient mapping using a histogram size of 6 bits and losslessly compressing the resulting signal with a pre-trained static Huffman encoder. Our results have been across validated using 3 datasets (Flint, Sabes and Brochier) and 3 new subjects suggesting consistent compression performance. The system has been implemented on a FPGA platform using 246 logic cells, consuming only 0.96 *µW* /channel and can accommodate more than 300 channels within 4kB RAM. All results and hardware designs are made publicly available, and researchers are free to select from them for their own system designs.

## Supporting information

Supplemental Material 2 - Results in excel format

## Author Contributions

O.W.S. and Z.Z. contributed equally to this work. Compression and behavioral decoding work was carried out by O.W.S., and hardware design and optimisation work was done by Z.Z. P.F. contributed the communication energy review. T.G.C. supervised the work, and directed and edited the manuscript.

## 6 Acknowledgements

Thank you to Robert D Flint, et al., Joseph E. O’Dohert et al. and Thomas Brochie et al. for making their neural and behavioral datasets publicly available [28, 32, 33]. O.W.S. is supported through an Physical Sciences Research Council (EPSRC) Doctoral Training Partnership (DTP) award (EP/N509486/1). P.F. was partly supported by the Engineering and EPSRC grant (EP/M020975/1, EP/R024642/1). This work was supported by the UK Dementia Research Institute which receives its funding from DRI Ltd, funded by the UK Medical Research Council, Alzheimer’s Society and Alzheimer’s Research UK.

---

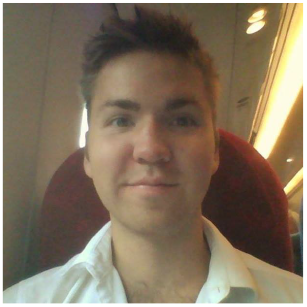

Oscar W. Savolainen became a Student Member of IEEE in 2018. He graduated with a B.Eng. degree from the University of Strathclyde, Glasgow, Scotland in 2018 and is currently pursuing his PhD at Imperial College London. His research interests include data compression, brain-machine interfaces, machine learning and time-frequency statistics. In particular, his main research focus is on investigating hardware-efficient means of compressing the Multi-Unit Activity and Entire Spiking Activity signals for wireless intracortical brain-machine interfaces. He has authored 8 papers, and during his undergraduate studies Mr. Savolainen was recipient of the Institute of Engineering Technology (IET) Power Academy scholarship, sponsored by National Grid.

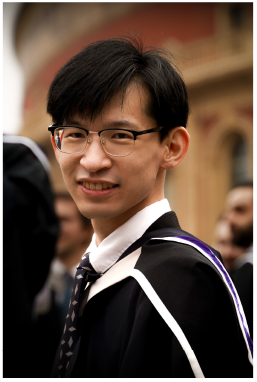

Zheng Zhang became a Student Member of IEEE in 2022. He received the B.Eng degree jointly from Beijing University of Technology (BJUT), China, and University College Dublin (UCD), Ireland, in 2018, and the M.Sc degree from Imperial College London in 2019. He is currently pursuing his Ph.D. at Imperial College London. His research interests are in hardware-efficient neural signal processing, brain-machine interfaces and artificial intelligence. His main research focus is on hardware-efficient real-time neural spike detection. He is the author of 6 papers, and during his undergraduate studies, he received the China National Scholarship and BJUT President Scholarship.

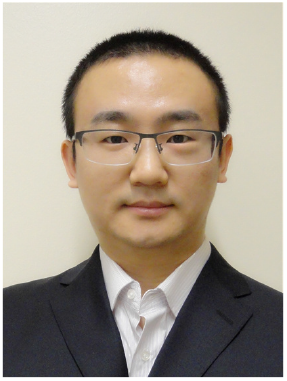

Peilong Feng (Member, IEEE) received the B.Eng degree in electrical engineering from the Henan Polytechnic University, China, in 2011, the first M.Sc degree in microelectronic systems design from University of Southampton, UK, in 2012, the second M.Sc degree in analogue and digital integrated circuit design from Imperial College London, UK, in 2015, the Ph.D. degree from the Imperial College London, in 2020. He is currently research associate at the Next Generation Neural Interfaces (NGNI) Laboratory. From 2012 to 2014, he worked as an electronic engineer in Shanghai Research Institute, China Coal Technology and Engineering group. His current research focuses on completely wireless infrastructure for distributed mm-sized neural implants and CMOS-memristor technologies.

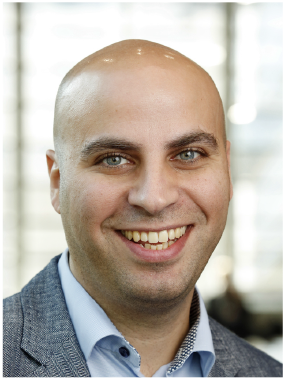

Timothy G. Constandinou (Senior Member, IEEE) received the B.Eng. and Ph.D. degrees in electronic engineering from Imperial College London, in 2001 and 2005, respectively. He is currently Professor of Bioelectronics at Imperial College London, Director of the Next Generation Neural Interfaces (NGNI) Laboratory, and Head of the Circuits & Systems Research Group. He is also a Group Leader within the UK Dementia Research Institute (DRI), Care Research and Technology Centre. His research interests are in microelectronics, biomedical microsystems, implantable medical devices, neural interfaces, brain-machine interfaces, research platforms, and remote sensing using ultra-wideband radar. He is a fellow of the Institution of Engineering Technology (IET), a Chartered Engineer, and a member of the Institute of Physics (IoP). He currently serves on the IEEE CAS Society, Sensory Systems and BioCAS Technical Committees. He was previously Associate Editor-in-Chief of IEEE Transactions on Biomedical Circuits and Systems (2020-2021) and served on the IEEE Circuits and Systems Society Board of Governors (2017–2019). He was the Technical Program Co-Chair of the IEEE BioCAS Conference, in 2010, 2011, and 2018, the General Chair of the BrainCAS 2016 and NeuroCAS 2018 Workshops.

