## Supplemental Material for "Hardware-Efficient Compression of Neural Multi-Unit Activity"

### Supplemental Material - Hardware-Efficient Compression of Neural Multi-Unit Activity

August 4, 2022

#### 1 Multiplexed Huffman encoder operation

An example scenario with two multiplexed Huffman encoders, encoding different channels of neural data, is given in Table 1. Channel 1 uses encoder $a$, channel 2 uses encoder $b$, etc. (Table 1 (c)). It can be observed that the multiplexed signal can be decoded if the symbol-channel-encoder couplings are known (Table 1 (d)).

Table 1: Demonstration of multiplexed Huffman encoding. (a) Huffman encoder $a$, trained on equal probabilities. (b) Huffman encoder $b$, trained on skewed probabilities. (c) Pairings of encoder to symbol sequence to channel. (d) Each symbol can be assigned to any encoder, and the sequence can be fully decoded if the pairings are known. This allows channels to use appropriate encoders from a selection.

**(a)**

| **Encoder a Huffman Table** | | |
|---|---|---|
| **Symbol** | **Frequency** | **Huffman Code** |
| 0 | 0.25 | 00 |
| 1 | 0.25 | 01 |
| 2 | 0.25 | 10 |
| 3 | 0.25 | 11 |

**(c)**

| **Symbol - Encoder - Channel pairings** | | |
|---|---|---|
| 1st Symbol | Encoder a | Channel 1 |
| 2nd Symbol | Encoder b | Channel 2 |
| 3rd symbol | Encoder b | Channel 3 |

**(b)**

| **Encoder b Huffman Table** | | |
|---|---|---|
| **Symbol** | **Frequency** | **Huffman Code** |
| 0 | 0.8 | 1 |
| 1 | 0.1 | 01 |
| 2 | 0.08 | 001 |
| 3 | 0.02 | 000 |

**(d)**

| **Encoding and Decoding Signal** | |
|---|---|
| Multiplexed signal (channel 1, channel 2, channel 3) | 3; 0; 1 |
| Encoded mult. signal | 11101 |
| Decoded mult. signal | From encoder a: 11 = 3;<br>From encoder b: 1 = 0;<br>From encoder b: 01 = 1; |

#### 2 Dataset details

To get a broad sample of MUA conditions, three datasets were used. The first was the Flint-Slutzky 2012 dataset [1]. It has 12 recordings taken across 5 days. Each recording consists of SUA data measured with a 96-channel Utah array in M1 in a non-human primate performing a free reaching task. The SUA data was obtained by taking broadband recording sampled at 30 kHz, highpass filtering them at 300 Hz, manually thresholding the resulting signal at an average of 5.2 standard deviations above the mean potential, and then manually sorting. In this work, MUA was derived from the SUA via collation of intra-channel spikes. In this work, the X and Y-axis cursor velocities, sampled at 100 Hz, were used as the observed behavioral data.

The second dataset was the Brochier et al. 2018 dataset [2]. This consisted of two recordings, one obtained from each of two non-human primates (referred to as L and N) from a 96-channel Utah array in both the M1 and the dorsal (PMd - subject L) or ventral (PMv - subject N) premotor cortex. The recording from subject L was approximately 1.5 hours long from a single session, and the recording from subject N was approximately an hour long, also from a single session. The broadband data was first filtered with a 1st order 0.3 Hz high pass filter (full-bandwidth mode) and a 3rd order 7.5 kHz Butterworth low pass filter. The SUA was then obtained via manual offline thresholding and sorting, from broadband data sampled at 30 kHz. For subject L, prior to thresholding, the data was highpassed at 250 Hz. Subject L had some noisy spikes that were not removed in this work. For subject N, prior to thresholding, the data was passband filtered between 0.25 and 5 kHz. The behavioural data consisted of labelled actions. As these were not continuous measurements, the behavioral data from Brochier et al. was not analysed in this work so as to keep the BDP metric consistent.

The third dataset was the O'Doherty Sabes lab 2017 (updated in 2020) dataset [3]. Two non-human primates (Indy and Loco) were recorded performing a free-reaching task. The neural recordings consisted of SUA from Utah arrays implanted in M1 (and for some sessions also in S1). As such there were either 96 or 192 channels per session, where some channels had no spiking activity and so were removed. 37 sessions spanning 10 months were recorded from Indy, and 10 sessions spanning a month from Loco. The hand and cursor X and Y positions were sampled at 250 Hz. The X and Y velocities were taken to be the observed behavioral data in this work, taken as the 1st derivative of position.

### 3 Data Compression analysis process

In this section, we determined the effect of various compression architectures on BR and therefore communication power. The role of the number of encoders, $S$, BP, and histogram memory size, which is used in assignment of channels to encoders, were investigated. However, firstly, not all static Huffman encoders will be useful for compressing MUA data at a given BP. As such, to identify the subset of interesting Huffman encoders, a Machine-Learning (ML) strategy was adopted. This is because the subset of interesting Huffman encoders, to be implemented on-implant, should be found in a way that is compatible with real-life application. I.e., the choice of the subset of Huffman encoders implemented on-implant should not be influenced by the data to be measured on-implant, as this is not known prior to implantation.

An approach was taken where all possible static Huffman encoders of length $S$ were considered. During each training-validation round, the set of Huffman encoders was reduced by removing the encoder that contributed the least to the compression. Eventually, only $u = 1$ encoder, i.e. the 'best encoder', was left, in which case no assignment or histogram were necessary and every channel used the same encoder. This is discussed further in the Supplemental Material, Section 3.3.1.

As required for a ML approach, a training/validation split was used. The training data was used to reduce the set of Huffman encoders. The validation data was used to represent neural data recorded on-implant, on which the quality of the assignment and compression was measured.

The training and validation data, from $A$, were split by taking the full recording from a channel and assigning it to either training or validation. The training and validation channels were randomly selected with a 50/50 split. The channels from the Flint and Sabes datasets were split separately, so that both were represented 50/50 in training and validation. All 960 Flint channels in $A$ were considered, and a random 2000 out of 4224 channels in the Sabes data in $A$ were considered. The Sabes data was limited so that the contribution of the Flint data was not overshadowed, and so prevented overfit to the Sabes data. It warrants mentioning that this training-validation split is distinct from that in the Supplementary Material, Section 4, that looked at the effect of BP and $S$ on BDP, although both splits concerned data from $A$.

#### 3.1 SCLV representation

We wanted to determine how the best $u$ ($u \in Z, 1 \leq u \leq h$) Huffman encoders, from the set of all possible $h$ Huffman encoders with $S$ fields, would perform in compressing MUA data from multiple channels where each channel was assigned to its ideal encoder. As such, all possible Huffman encoders of length $S$ were produced. For example, for $S = 3$, all possible Huffman encoder codeword combination are given in Table 2 (a).

There are many redundant configurations of Huffman encoders, assuming the order of the symbols in unimportant. The only constraint is that, in a Huffman encoder, the shortest codeword should represent the most likely symbol in the data, the 2nd shortest should match the 2nd most likely, etc. As such, the Huffman encoders were reduced down to a Sorted Codeword Length Vector (SCLV) representation, which consisted of the ascending sorted vector of codeword lengths. This process is shown in Table 2 (a-c). In the case of $S = 3$, it can also be observed in Table 2 (c) that all $h = 6$ encoders reduce down to a single SCLV. Taking the dot product of the SCLV and Sorted Histogram (SH), where the histogram is sorted in descending order, gives the length of the data, in bits, after compression by a Huffman encoder with the same SCLV as shown in Table 3 (c). Dividing the dot product by the number of samples gives $avlen$ from Eq. 2 in the main manuscript, where dividing further by BP gives the BR. This enables

Table 2: Demonstrating a non-redundant representation of Huffman encoder codeword lengths, the SCLV. (a) All Huffman code combinations for S = 3; (b) Vectors of the lengths of the Huffman codes, with copies removed. (c) Sorted Codeword Length Vectors, with copies removed.

**(a)**

| **All Huffman Encoders of Length S = 3** | | | |
|---|---|---|---|
| **Huffman code combination** | **1st CW*** | **2nd CW** | **3rd CW** |
| **1** | 0 | 10 | 11 |
| **2** | 0 | 11 | 10 |
| **3** | 00 | 1 | 01 |
| **4** | 01 | 00 | 1 |
| **5** | 11 | 0 | 10 |
| **6** | 00 | 01 | 1 |

***CW = Codeword**

**(b)**

| **Non-redundant Codeword Length Vectors, S = 3** | | | |
|---|---|---|---|
| **Huffman code length combination, key** | **1st CW length** | **2nd CW length** | **3rd CW length** |
| **1** | 1 | 2 | 2 |
| **2** | 2 | 1 | 2 |
| **3** | 2 | 2 | 1 |

**(c)**

| **Non-redundant Sorted Codeword Length Vector (SCLV)** | | | |
|---|---|---|---|
| **SCLV combination** | **1st CW length** | **2nd CW length** | **3rd CW length** |
| **1** | 1 | 2 | 2 |

us to assign encoders to channels based on which encoder gives the smallest BR (Table 3 (d)). It warrants mentioning that, to actually compress the data, a Huffman encoder that matches the SCLV is required. However the BR can be obtained from the SCLV via its dot product with the SH.

Based on the BDP vs. $S$ results that will be shown in Supp. Mat. Section 5, and our knowledge of the hardware costs of large $S$ values that will be shown in Supp. Mat. Section 7, we opted for the full integrated results to only examine $S$ values between 2 and 9 for our set of BPs. This allowed relatively good BDP values, as discussed later, while minimising the resources. Therefore, for each integer value of $S\,\{S \in \mathbb{Z}, 2 \leq S \leq 9\}$, the full set of SCLVs was produced. The details are given in the next section.

#### 3.2 Producing the Full Set of Huffman Sorted Codeword Length Vectors

For each integer value of $S\,\{S \in \mathbb{Z}, 2 \leq S \leq 9\}$, the full set of SCLVs was produced by creating every non-redundant combination of probability vectors $p$ of length $S$, at a given discrete resolution $q$. Each probability value $p[k]$, with $k\{k \in \mathbb{Z}, 1 \leq k \leq S\}$, was equal to $q \times j$, where $q$ was the discrete increment and $j \in \mathbb{Z}^+$ so that $0 \leq p[k] = q \times j \leq 1$. $q$ was set to a low value of 0.1, and each non-redundant probability vector $p$ of length $S$ was created.

Each $p$ was then normalised to $p'$ so all of its $k$ elements summed to 1. A Huffman encoder was then trained on $p'$, where $p'$ was used to represented the frequencies of arbitrary symbols. Each Huffman encoder was then reduced down to the SCLV representation, and the non-redundant SCLVs for each $S$ stored. With small enough $q$, every possible Huffman encoder, and so every possible SCLV, is generated.

#### 3.3 ML process

##### 3.3.1 Training

The system was trained by assigning the SCLVs to the training channels based on which SCLV gave the best BR for each channel. Multiple channels could be assigned the same SCLV, and all of the data in each channel was used.

To identify interesting SCLVs, i.e. train the system, the following strategy was used. Firstly, each training channel's SH was obtained. Secondly, one by one, with replacement, each SCLV was removed. The average BR across channels was then measured after each channel was assigned to its best remaining SCLV, e.g. as in Table 3. Channels could be assigned to any SCLV, except the missing one. After the total BR had been obtained for each missing SCLV, the SCLV that was found to be the least effective SCLV for compressing the training data, in concert with the other SCLVs, was removed. As such, at the end of the training round, one SCLV had been removed. This process is represented in Fig. 1 (b).

Table 3: Taking the dot product between the SCLVs (a) and example Channel Sorted Histograms (b) to obtain the size of the compressed data blocks (c). From (c), we can determine the best SCLV-channel pairs. In (d), the channel-average length $avlen$ in bits of the encoded symbols after each channel is compressed using its ideal encoder is obtained, with $L = 1000$.

(a)

| SCLVs | | | | | |
|---|---|---|---|---|---|
| **SCLV \Symbol** | **0** | **1** | **2** | **3** | **4** |
| **1** | 1 | 2 | 3 | 4 | 4 |
| **2** | 2 | 2 | 2 | 3 | 3 |
| **3** | 1 | 3 | 3 | 3 | 3 |

(b)

| Channel Sorted Histograms (SH) | | | | |
|---|---|---|---|---|
| **Symbol \Chan.** | **1** | **2** | **3** | **4** |
| **0** | 650 | 500 | 300 | 950 |
| **1** | 200 | 200 | 300 | 5 |
| **2** | 100 | 120 | 200 | 31 |
| **3** | 50 | 100 | 100 | 7 |
| **4** | 0 | 80 | 100 | 7 |

(c)

| Dot Product of Channel SHs and SCLVs | | | | |
|---|---|---|---|---|
| **SCLV \Chan.** | **1** | **2** | **3** | **4** |
| **1** | **1550** | **1980** | 2300 | 1109 |
| **2** | 2050 | 2180 | **2200** | 2014 |
| **3** | 1700 | 2000 | 2400 | **1100** |
| **Best SCLV** | **1** | **1** | **2** | **3** |

(d)

$$avlen = \frac{1}{n}\Sigma_{i=1}^{n}\frac{min(dotprod[:,i])}{L}$$

$$avlen = \frac{1}{4}\frac{1550 + 1980 + 2200 + 1100}{1000} = 1.7075 \text{ bits}$$

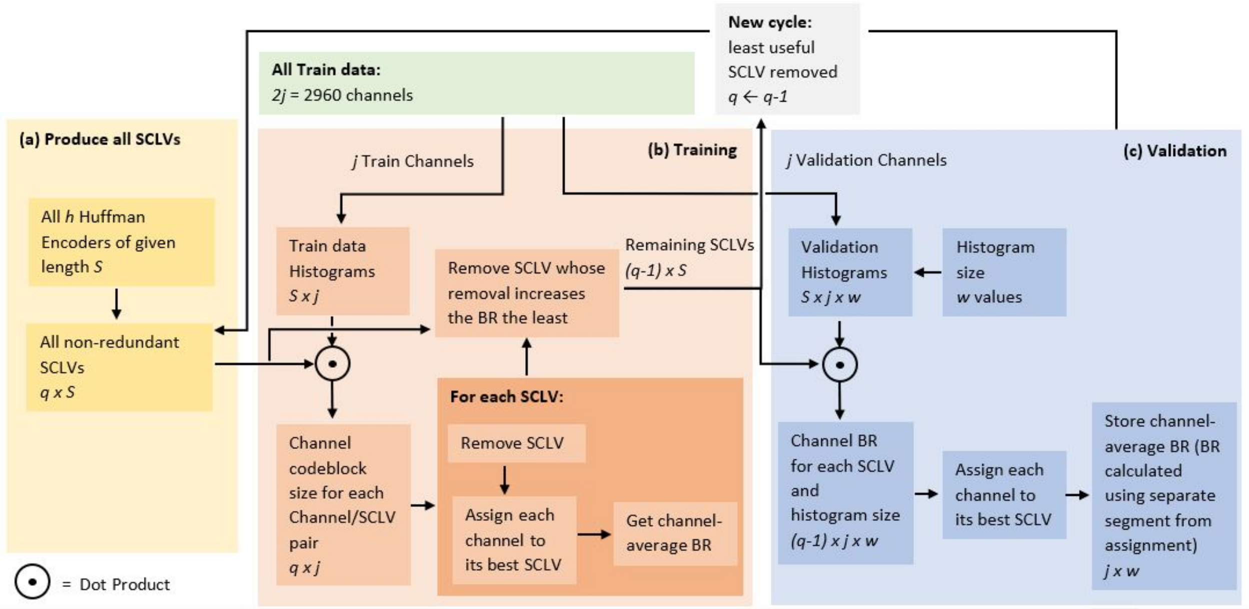


Figure 1: Encoder-selection ML algorithm. Italics represent data dimensions. First, all possible SCLVs are produced (as shown in (a)). Then, each round selects an increasingly smaller subset of SCLVs based on which give the best compression in the training set (as represented in (b)). At each round, i.e. with different numbers of SCLVs, the compression is tested on the validation dataset for different on-implant histogram sizes (as represented in (c)).

###### 3.3.2 Validation

Between each training round, validation was performed. The purpose of validation was to measure the BR when each 'on-implant' channel was assigned to its ideal encoder from the selection of encoders available during the current training round. In other words, it was tested what the on-implant BR would be given the selection of encoders found during that training round. The method of assigning on-implant MUA channels to encoders should be realistic to implement in hardware. In this work, the assignment was done by using a segment from the beginning of each on-implant channel recording to produce a sample histogram. Each channel was then assigned to an encoder based on which encoder gave its histogram the smallest BR. This is represented in Fig. 1 (c).

How much of the beginning of the 'on-implant' validation recording was used to calculate the sample histogram depended on the size of the histogram. The considered histogram sizes were $S \times 2^d - 1$ samples, where $d \in [2, 3, 4, ..., 9, 10]$. Each histogram bin, of which there were $S$, was given a size of $2^d - 1$ maximum samples, i.e. of $d$ bits. Once $2^d - 1$ samples had been measured across all bins, the histogram growth ended, the histogram was sorted, and the channel was assigned to an encoder.

After assignment, the resulting BR was calculated. This was done for each channel by taking the rest of the validation data, that which had not been used for assignment, and calculating its BR after compression. In particular, only a segment of the remaining data was used, where the segment was always equal in length to half of the total recording length. This was so that the amount of compressed data was the same across all histogram sizes. This is shown in Fig. 2, where the beginning of the recording is used for assignment, and a segment of the rest for calculating the BR. The average BR across validation channels was then stored for each histogram size.

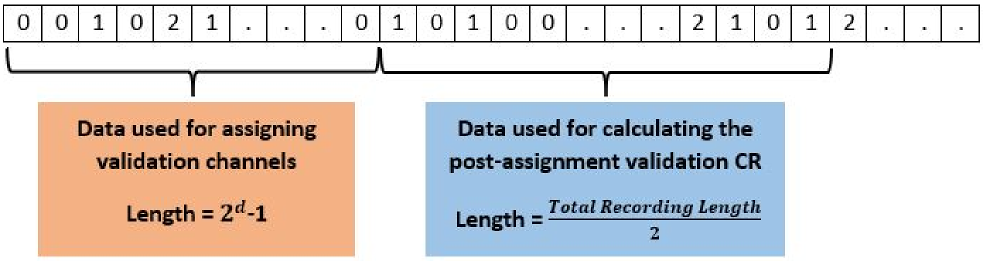


Figure 2: Single-channel split of assignment (for sample histogram) and to-be-compressed validation data, based on histogram memory size $d$ and total recording length.

We then moved to the next training-validation round. At each round, the least useful SCLV was removed based on the training data. The validation channels were then assigned to the remaining SCLVs, and the average BR for each histogram memory size stored. The process continued until only 1 encoder (i.e. the best encoder for the training data) was left, where all validation channels were automatically assigned to the last encoder. As such, for each round (i.e. number of non-redundant SCLVs), the mean BR, ideal set of encoders, and effect of histogram size on BR were all determined. 30 different cross-validation iterations of this process were run in parallel, with different training-validation channel splits, and the results were combined.

It is important to mention that, for any channel, the compressed on-implant data should be sorted the same way its histogram was sorted. For example, if a firing rate of 3 MUA events per bin is found to be the most common firing rate in the sample histogram, the occurrence of 3 MUA events in a bin in the remaining to-be-compressed data should be given the shortest codeword during compression. This is because the sorted ordering in the compressed data has to be determined somehow, and so is approximated using the sample histogram for each channel. As such, the histogram serves to not only get an idea of the shape of the channel's histogram, used to find the ideal encoder for compression, but also to find the sorting order of the to-be-compressed data, as the most common firing rates should be assigned the shortest codewords within each encoder. We refer to this assignment of to-be-compressed firing rates to encoder fields, based on the sample histogram, as mapping (e.g. mapping the symbols to the encoder fields). As such, the size of the histogram affects the quality of both the assignment and the mapping, both which impact the BR. The sorting and mapping implementation is discussed further in Section 6.

### 4 Parameter optimisation during determining the effect of $S$ and BP on BDP

The only data that was observed was the training data $A$ specified in Section 2.1.2 of the main manuscript. Each recording in $A$ was split into training and testing, using a 90-10% split. The training data was then split into sub-training and validation using 5-fold cross-validation. The WCF hyper-parameters and the pre-processing parameters were then optimised on the folds. The optimised decoder hyper-parameters included the WCF degree, the alpha parameter, and the number of taps i.e. timesteps used in the WCF. $l_2$ (i.e. ridge or Tikhonov) regularisation was used by default. The optimised pre-processing parameters included the lag between the neural and behavioral data, and the width of the causal moving average window used to smooth the neural data. The optimised parameter values are given in Table 4.

Table 4: Optimised parameters and hyper-parameters for the behavioral decoders.

| | |
|---|---|
| **Timesteps** | 5, 10, 15 |
| **Kinematic to neural lag (s)** | 0, 0.02, 0.04 |
| **Moving average window length (s)** | 0, 0.05, 0.1, 0.2 |
| **Alpha** | 0, 1e-4, 1e-2 |
| **CWT Degree** | 2, 3, 4 |

The best-performing parameters across the 5 folds for each $S$ and BP were then used as the parameters for the decoder tested on the testing data, giving the BDP as a function of $S$ and BP. Here, for testing the effect of $S$ on BDP, BPs of 1, 5, 10, 20, 50 and 100 ms were investigated.

### 5 BDP Results

#### 5.1 BDP as a function of BP and $S$ - Train data $A$

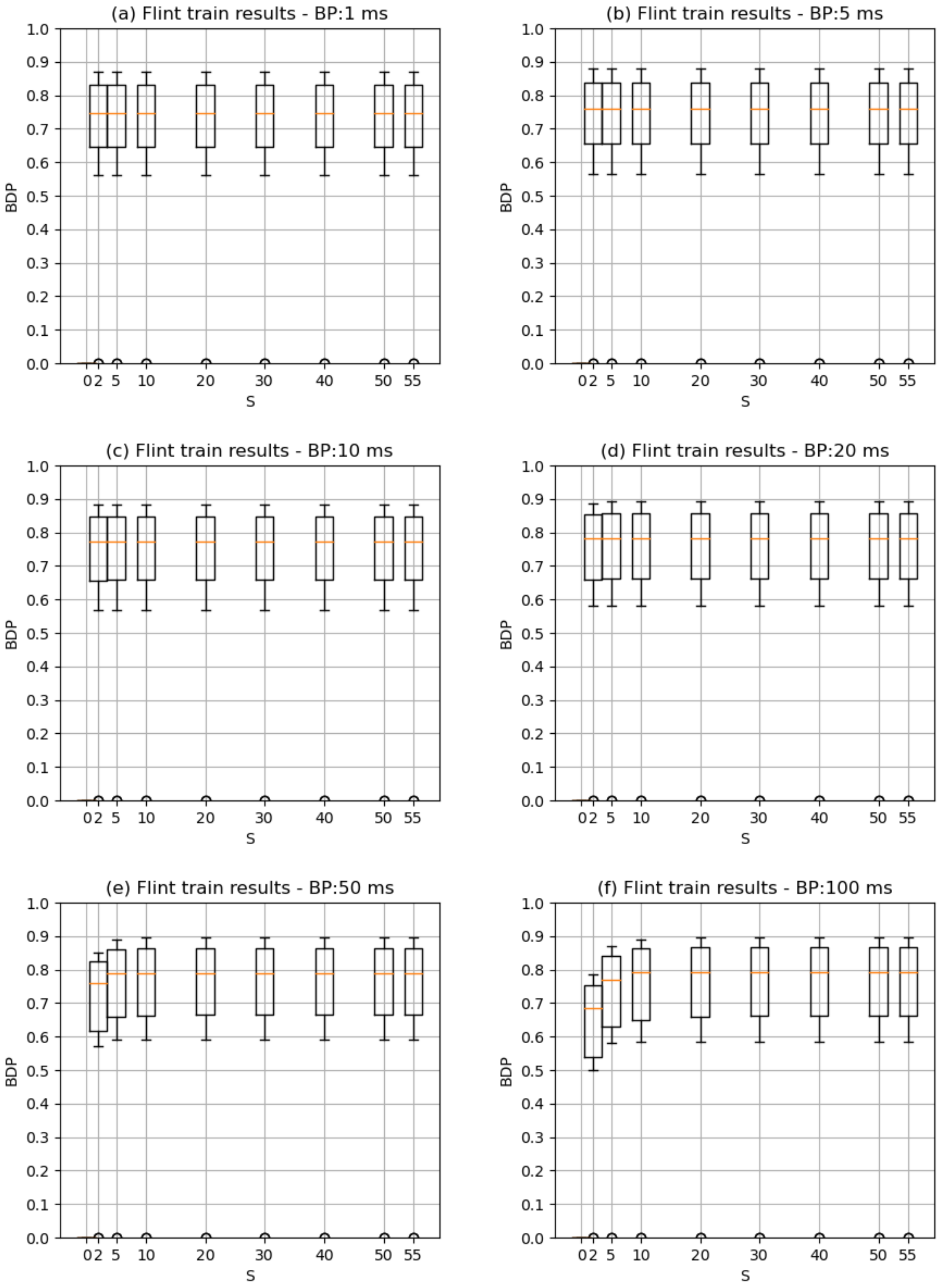


Figure 3: Behavioral decoding performance (BDP) as a function of BP and $S$ for the Flint data in $A$. Each S/BP combination was parameter optimised on 5-fold CV.

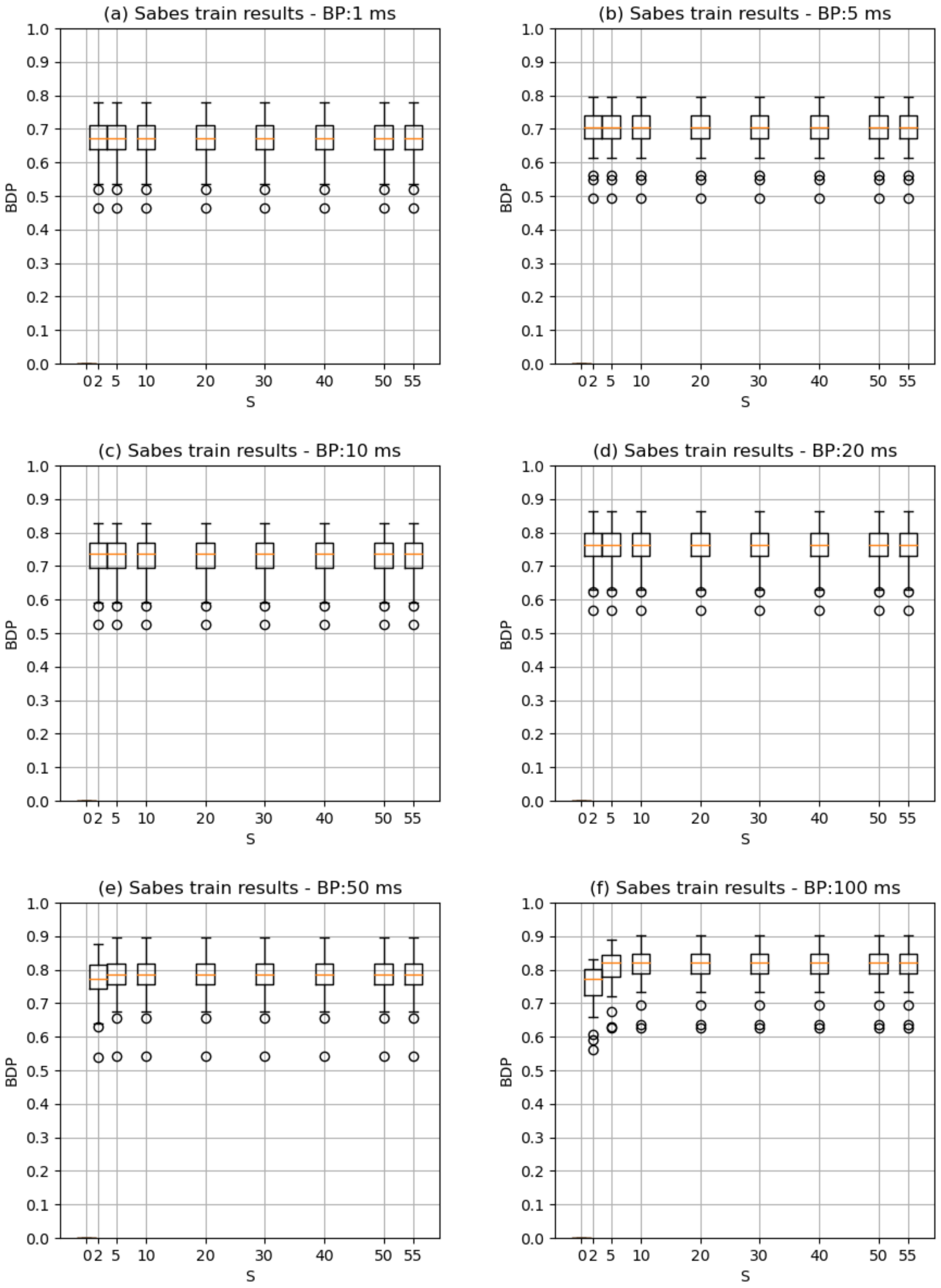


Figure 4: Behavioral decoding performance (BDP) as a function of BP and $S$ for the Sabes data in $A$. Each S/BP combination was parameter optimised on 5-fold CV.

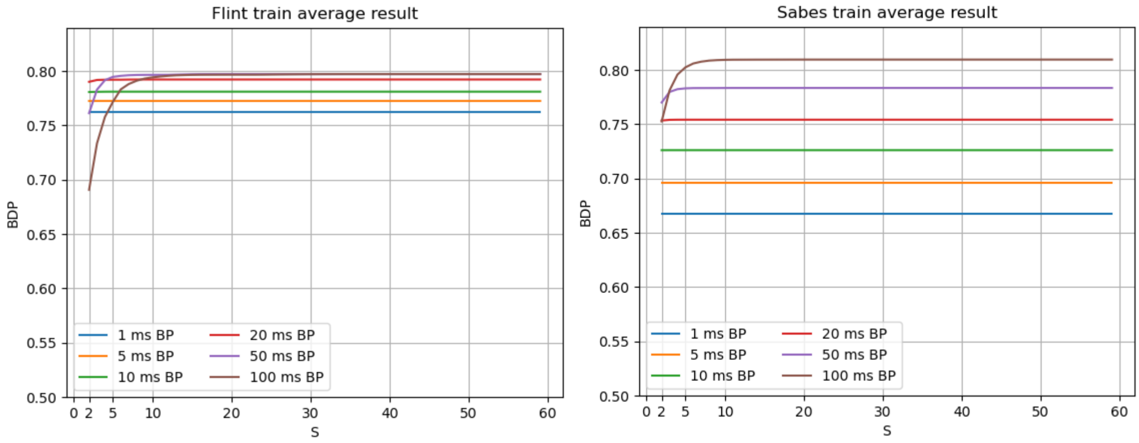


Figure 5: Behavioral decoding performance (BDP) for Flint and Sabes training data as a function of BP and $S$, averaged across the recordings.

From Fig. 5, it warrants mentioning that BDP increases with BP, assuming the effect of increasing $S$ has saturated to that of $S = max(x)$ (where the line stabilizes horizontally for each BP). However, the value at which the effect of $S$ saturates increases for each BP. Interestingly, a lower BP can have a better BDP than a higher BP at the same $S$, since no information is being saturated for the lower BP. However, even low $S$ values perform quite well relative to the peak values, even for a BP of 100 ms. It is also interesting that these results are opposite to those in [4], where BDP decreased linearly as a function of BP. However, [4] used LSTM decoders whereas this work uses WCFs. It isn't surprising that deep learning algorithms may be able to find more information because at higher temporal resolution than simple decoders like the WCF. Also, [4] looked at the average effect of BP across different signal processing parameters, whereas this work looked at the effects when using fully optimised system parameters. It also warrants mention that, even if one generally gains BDP at a higher BP, one loses on temporal resolution of the decoded output. Furthermore, one can always bin the data to a higher BP prior to decoding if one wishes to gain a better BDP, as higher BPs are simply a lossy representations of lower BPs.

#### 5.2 BDP as a function of BP and $S$ - Test data $B$

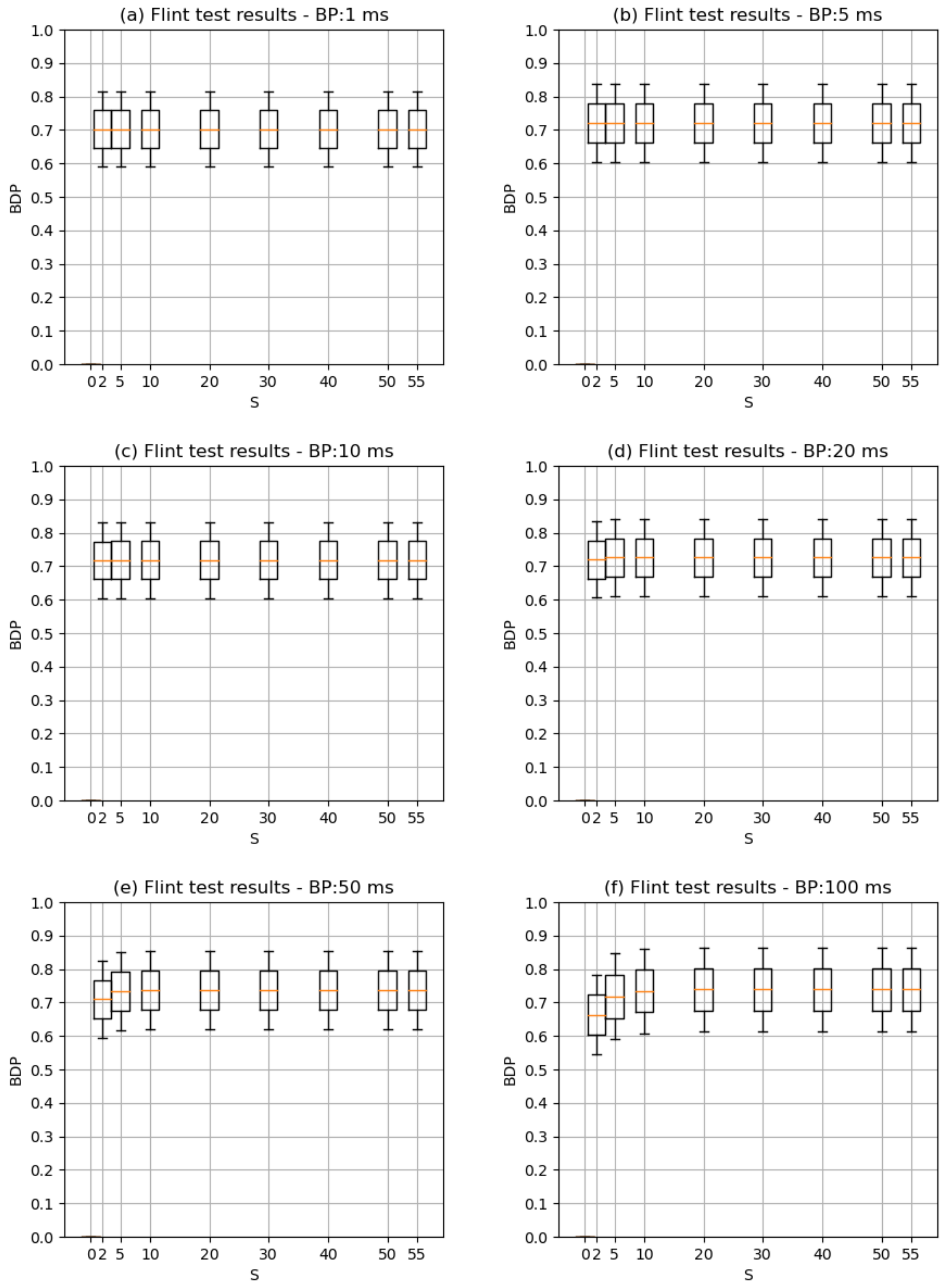


Figure 6: Behavioral decoding performance (BDP) as a function of BP and $S$ for the Flint data in $A$. Each S/BP combination was parameter optimised on 5-fold CV.

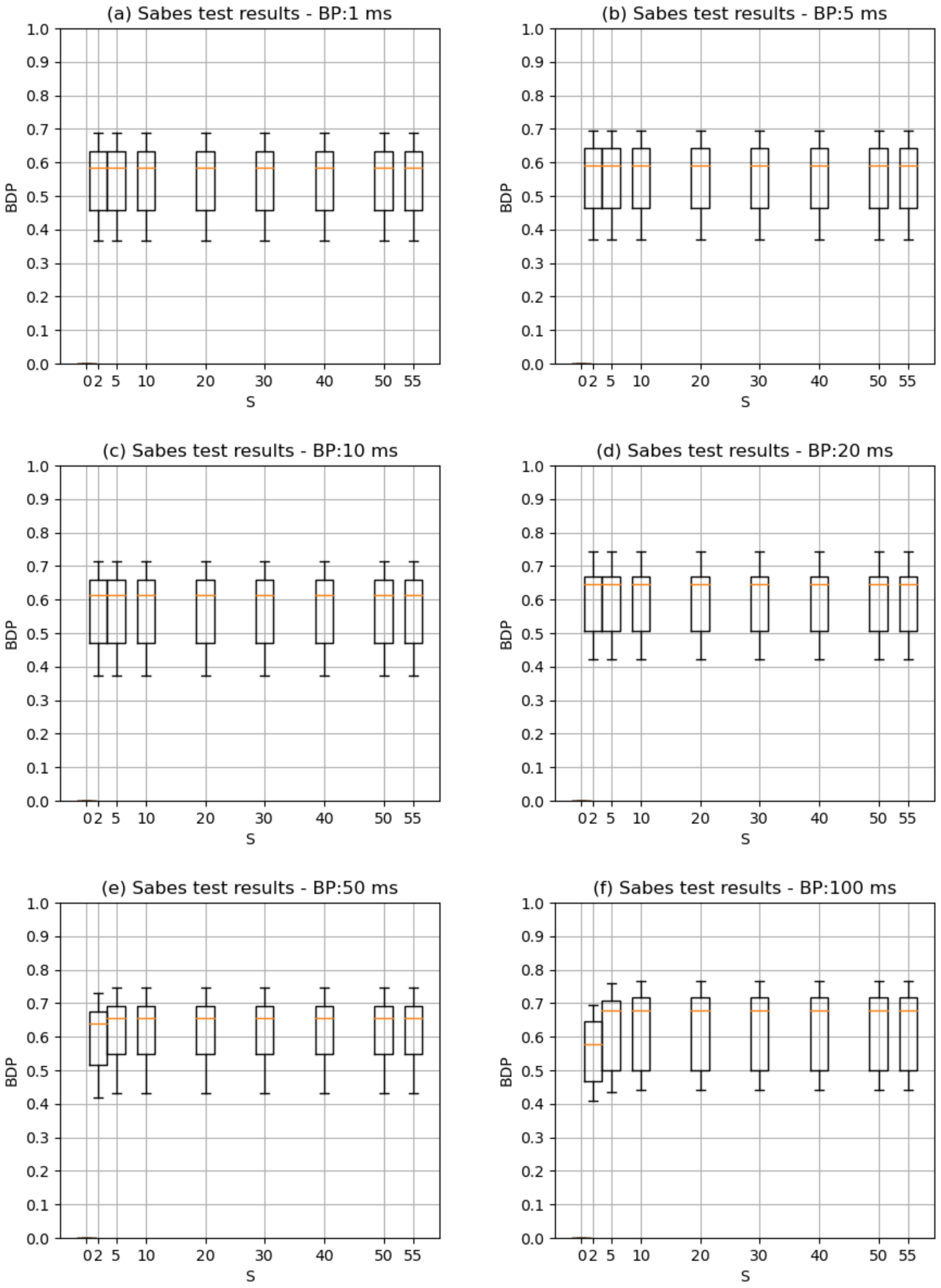


Figure 7: Behavioral decoding performance (BDP) as a function of BP and $S$ for the Sabes data in $A$. Each S/BP combination was parameter optimised on 5-fold CV.

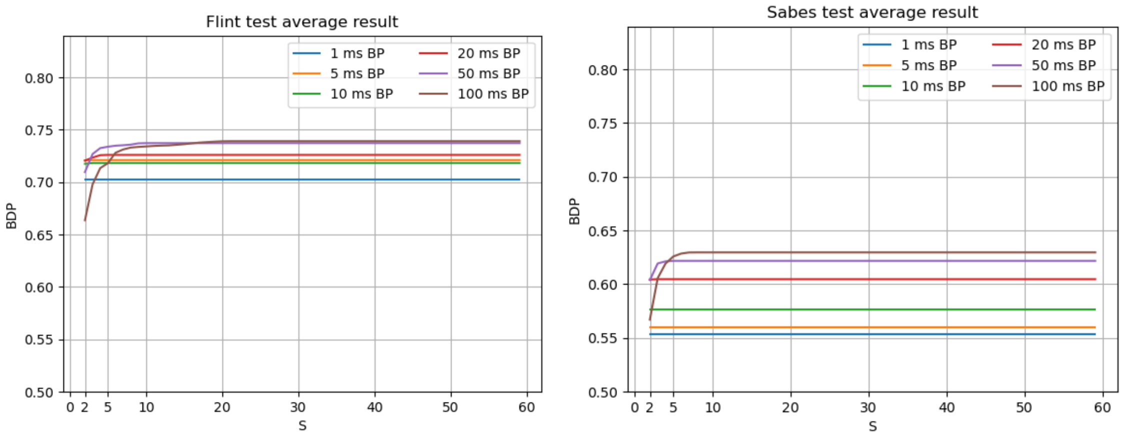


Figure 8: Behavioral decoding performance (BDP) for Flint and Sabes testing data as a function of BP and $S$, averaged across the recordings.

From Fig. 8, it can be seen that the BP/$S$ relationship to BDP holds much the same in the test data $B$. However, the BDP values are generally significantly lower in the training data (Fig. 5). The same parameter optimisation was performed, and so either the range of optimised parameters is non-ideal, the test recordings have less behavioral data in them, or they are more noisy.

### 6 FPGA realisation

We simulated the presented architectures on an FPGA target, Lattice ice40LP, in order to assess the overhead on power and resources brought by compression so as to guide our configuration selection. Such a low-power, high-performance FPGA with 40nm technology and small BGA package is ideal for the thinnest devices like implantable BMIs. All programs are written in Verilog, simulated on Mentor Modelsim Lattice Edition and synthesised with iCECube 2020.12.

Lattice ice40LP1K is an ultra-low-power FPGA board with 1280 logic cells and sixteen 4kbit memory blocks (bRAMs). The architecture of the FPGA implementation is shown in Fig. 9, including the Binner, Histogram counter, Sorter, Mapper, Encoder selector, Encoders and Memory. Referring to Table 7 and Fig. 5 in the main manuscript, different configurations can be achieved by bypassing some of the components. In the 'one encoder' version, there is only one encoder in Encoders and therefore no need for the Encoder selector module. In the 'Without Mapping' configuration, we assume the events in the histogram are already sorted and the sorter and mapper are bypassed. The *Spike rate freq* is connected with the *Sorted freq* in the encoder selector, and the *New spike rate* is connected with the *Mapped spike rate* in Encoders. Similarly, in the 'Only Binning' version, only the Binner module is implemented. A detailed breakdown of Fig. 9 is given below.

#### 6.1 Binner

A timer and a recurrent spike rate counter is used for the Binner, which counts the number of in-coming spike numbers in a given BP. The multiplexer (MUX) after the spike rate counter is used to clip the spike rate within the preset range and reset the spike rates stored in memory when a BP is over.

#### 6.2 Histogram

The Histogram is implemented using a Finite State Machine. The state transition diagram is shown in Fig. 9 (B). It accumulates the number of MUA events according to the *Bin finished* and new spike rate signals. When the histogram overflows, a finish signal is issued and the histogram is emptied for the next channel. As the maximum spike rate is clipped at $S - 1$, the number of registers required for storing the different MUA frequencies is highly reduced. The index of the maximum value in the MUA histogram is recorded in the histogram counter, to be used in the Sorter.

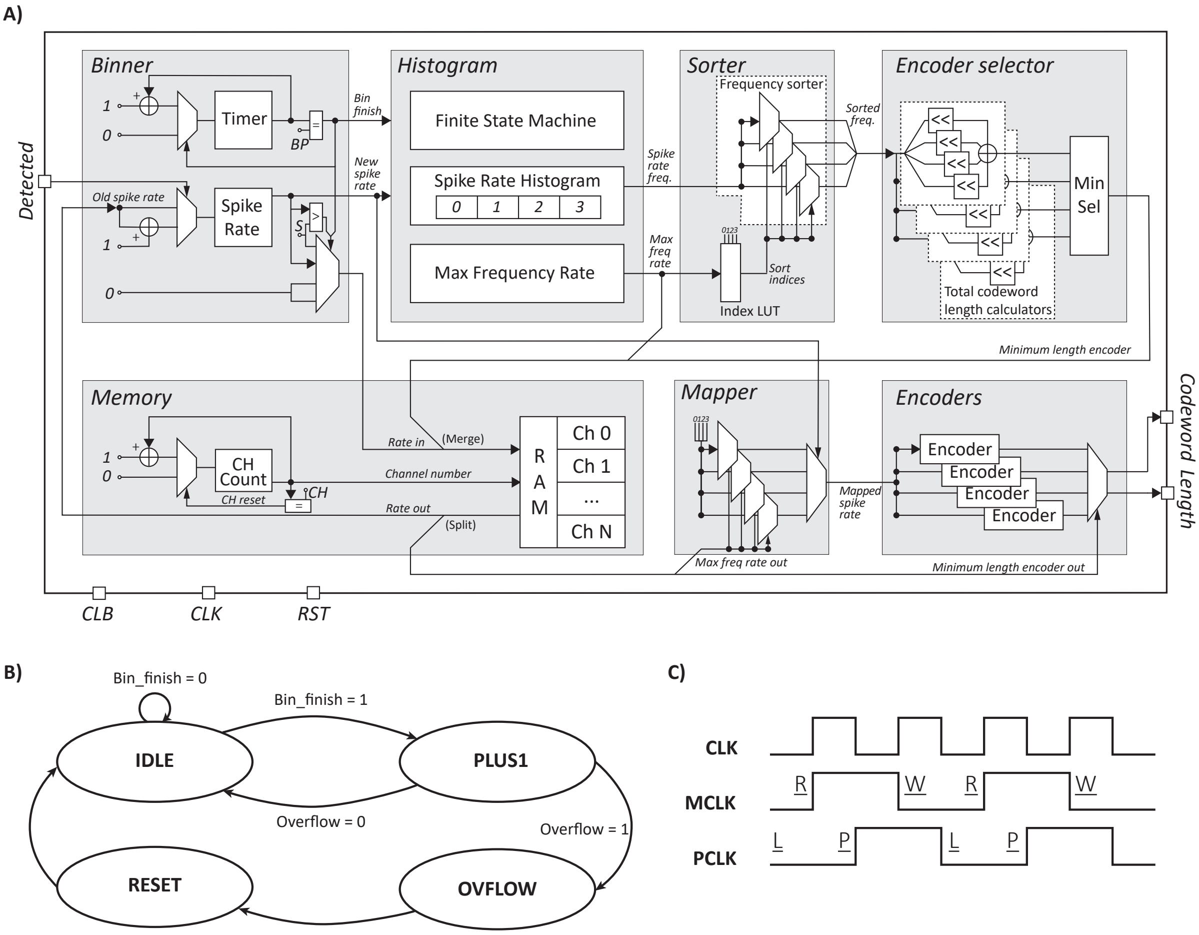


Figure 9: A). The FPGA implementation includes Binner, Histogram, Sorter, Mapper, Encoder selector, Encoders and Memory. For conciseness, several circuits are not shown: Clock generator for memory clock and processing clock, reset and re-calibration logic, clip for *New spike rate* and MUXs selecting *Minimum length encoder out*, *Sorted indices out* as the input to the RAM after calibration. B). State transition diagram of the finate state machine in Histogram module. C). Clock timing diagram of the system clock (CLK), Memory clock (MCLK) and Processing clock (PCLK). MCLK drives the RAM, it is read at the posedge (R) and written at the negedge (W). PCLK trigers the Binner and Histogram at its posedge (P). It also synchronise the detected signal at its negedge (L). The update of channel number is also happened at the negedge of PCLK (L).

#### 6.3 Sorter and mapper

Sorting the spike rate frequency in order can be resource-hungry or time-consuming in hardware. The resources used for implementing a sorting algorithm such as merge sort or quick sort can overwhelm the whole system. For sorting MUA histograms, we can take advantage of the fact that the MUA histogram, even if it does not follow a decaying exponential, is almost always expected to follow a unimodal peak distribution. As such, the Sorted Histogram (SH) can be easily estimated by setting the index 0 at the index of the histogram maximum, and the index number will increment by iterating on both sides of the histogram peak. For example, values to the left of the peak will take odd index numbers of 1, 3, 5, etc., while values to the right of the peak will take even numbers. When indices can no longer be assigned on one side, the rest are assigned serially to the other side. An example is illustrated in Fig. 10 A. Using such an estimation, we can reduce both the sorting space and time complexity to $O(n)$.

From a hardware perspective, this estimated sort algorithm can be implemented with a finite state machine (FSM). However, in Fig. 9, we show a combinatorial implementation. As the estimated sort order is only affected by the most frequent spike rate index, we can easily create a LUT that defines the sorting order based on the measured maximum index. The spike rate Mapper can also be implemented using an identical LUT. A demo of the sorted indices LUT when $S = 5$ is given in Fig. 10. We also implemented two other sort algorithms: a swapping sort and estimation

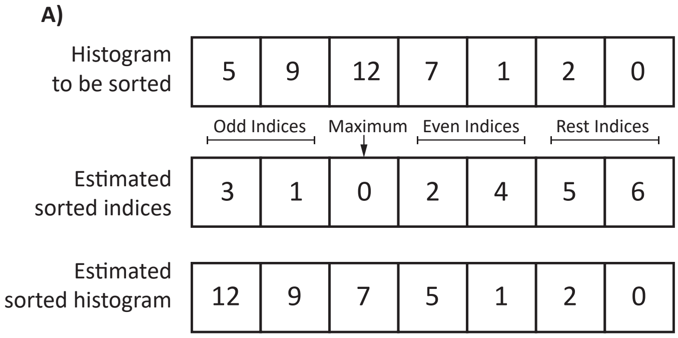


B)

| Max/Index | 0 | 1 | 2 | 3 | 4 |
|---|---|---|---|---|---|
| 0 | 0 | 1 | 2 | 3 | 4 |
| 1 | 1 | 0 | 2 | 3 | 4 |
| 2 | 3 | 1 | 0 | 2 | 4 |
| 3 | 4 | 3 | 1 | 0 | 2 |
| 4 | 4 | 3 | 2 | 1 | 0 |

Figure 10: A). A demo for the estimated sort algorithm. Index 0 of the sorted histogram will be the maximum of the unsorted histogram. Odd indices will be placed on the left while the even indices will be placed on the right. The rest of the indices will be placed serially on the unsorted side. B). A demo of the sorted indices when $S = 5$. Max indicates the index of the peak value in the histogram, and Index is the histogram field. Based on the peak value, the indices in the table are given to the histogram fields.

sort with sequential logic, which are given in Supplemental Material Section 5.

#### 6.4 Encoder Selector

The encoder selector assigns encoders to channels. For the given channel, it selects the encoder that gives the minimum encoded data length, determined by taking the dot product of the SH with the SCLVs. The SCLVs are stored on-implant for each encoder. Since multiplications are resource-hungry in hardware, we replace the multiplications with bit shifts. The number of bits to shift for different encoders and codewords are stored on board.

#### 6.5 Encoders

A Huffman encoder, in hardware, is a big LUT. Multiple encoders have been implemented on board. Different encoders are selected according to the encoder selector. The selected set of Huffman codewords and codeword length are then set as the outputs of this channel.

#### 6.6 Memory unit

The Memory unit consists firstly of a channel counter that counts the processed channels to schedule the data flow among channels. Secondly it consists of RAM that stores each channels' current MUA FR, their sample histogram largest value's index during calibration (used for mapping during regular encoding operation) and their assigned encoders. Instead of using registers for each channel, utilising RAM increases the scalability, making it possible to upscale to thousands of channels within only hundreds of logic cells. However, the clock speed needs to be doubled to maintain the same data throughput. As the resources are highly constrained in lattice iCE40LP, the RAM implementation is preferred over using registers.

Fig. 9 (c) shows the timing of the clocks. CLK is the system clock and a clock generator is used to generate MCLK for memory and PCLK for different processing units. The detection signal will be loaded at the negedge of the PCLK. The RAM will provide the stored parameters for different processing units at MCLK posedge. All processing happens on PCLK posedge and the results are stored back at the MCLK negedge. These modules work together to compress the MUA data of each channel. The histogram counter, sorter and encoder selector are used at the start of implant operation for encoder selection as a calibration process. During the calibration, for each channel, the sample histogram largest value's index and encoder assignment will be determined and stored into RAM. One can also periodically recalibrate each channel when the brain environment changes, or at some defined interval. After the calibration, the spike detection signal of different channels will flow through the binner, mapper, encoder and RAM interchangeably for compression.

#### 6.7 Hardware implementation of the sort algorithms

Utilising the nature of the spike rate, we can customise sort operations to reduce the hardware cost. Besides the combinatorial implementation introduced in the main context. We here proposed other two different sort algorithms, we named it a real-time swap sort and the other is an estimated sort with sequential logic.

##### 6.7.1 Swapping sort

The real-time swap sort the spike rate count histogram and also the order of spike rate (count lags) in real-time during histogram accumulation. It is therefore balances the computation load temporally. Based on the fact that every bin period the histogram counter reads the output spike rate from the binner, the spike histogram count can only be increased by 1. Therefore, swapping the updated spike rate count with the last value smaller than it can ensure the spike count histogram is in order.

In one iteration, the updated value only compare itself with the spike rate counts equal to the value before it is updated. The complexity is still $O(n)$ theoretically if the spike rate is random. However, as the actually spike rate count is expected to follow a unimodal peak distribution, the number of comparisons is much lower than $O(n)$ in practice.

##### 6.7.2 Estimation sort with sequential logic

Three important registers and two register banks have been used: *Max* stores the most frequent firing rate, which is obtained in histogram count; *Start* stores the current location of the left of the most frequency firing rate being sorted; *End* stores the current location on the right of the most frequency firing rate being sorted; *Freq* is a bank of S registers that store the estimated sorted firing rate frequencies; *Rate* a bank of S registers that store the original lags (firing rate) of the frequencies before sorting.

The sorter stays at *IDLE* when counting the firing rate histogram. After counting finishing, the sort enters *LOAD* state, in which it will load the frequency of each firing rate from histogram into *Freq*, stores 0 to S-1 in to *Rate* and set the current *Start* and *End* indices according to *Max*. When *Max* is 0, *Start* = 0. When *Max* is S - 1 (the largest possible firing rate), *Start* = S - 1, *End* = S. In other cases, *Start* = *Max* - 1, *End* = *Max* + 1. It will also store *Max* and its frequency at the first index of *Rate* and *Max*. If neither *End* = S nor *Start* = 0, the sorter will enter the *SORT* state, where *Start* and *End* will be stored in the next even index and odd index in *Rate*, the frequencies will also be stored in *Freq* accordingly. After that, *Start* will be reduced by one and *End* will be increased by one. If *End* = S, the sorter will enter *RSRT* state. In that state, the right side of the *Max* has been sorted, and only the left side, i.e. *Start* side will be processed. When *Start* becomes zero, sorting is finished. Note that, if *Start* is decreased to zero (the left side sort finishing), before *End* is increased S, the sorter will also enter *IDLE* state, as the rest on the right is already sorted.

Such an estimated sort, in the worst case when the maximum is the last value in the histogram, will take S clocks to get the result. However, the maximum is normally only appeared in the left half and it takes less than S/2 clocks after histogram count.

### 7 Hardware results

#### 7.1 Resource usage

Resources usage reflects the area occupation of the implementation. The full results are given in Supplemental Material 2 as an excel spreadsheet, and the same is available on the Github at [5]. The histogram and encoder selector are two resource-hungry modules. The amount of resources required by the histogram is mostly dependent on increased histogram size, but the $S$ values also have a significant but smaller impact. The resource usage of the encoder selector can exceed that of the histogram when $S$ is larger than 7 because of the dot product (implemented with bit shifting) between two vectors with length $S$. Alternatively, all possible multiplication results could be pre-calculated and stored in a RAM. The selector logic would be simplified, however this would sacrifice the processing speed as the results would need to be fetched from RAM one by one and summed together. As a result, the calibration time would be increased. Such an alternative approach could be useful if the limitation of the resources is extreme or the configuration requires a large $S$, histogram size or number of encoders. It would be less effective when these values are small. Aiming at finding the most compact compression scheme, we opted to use the bit-shift implementation discussed in the Sup. Mat., Section 6.

For the sorter, we have compared three different approaches: Swapping sort, and estimation sort with sequential and combinatorial implementations. A summary of the resources needed for the three implementations is given in Table 5.

One can notice that compared to the Swapping sort, the sequential estimation sort reduced the required resources by half, which makes it possible to do on-implant sorting with limited resources available. More noticeable, when $S$ = 5, histogram size = 4, the resources of the combinatorial implementation is only one-third of the sequential

Table 5: Logic cells were used in two different settings for three sort implementations.

| | Swapping sort | Estimation sort - Seq | Estimation sort - Comb |
|---|---|---|---|
| S = 5, hist size = 4 | 480 | 239 | 75 |
| S = 9, hist size = 6 | 682 | 355 | 260 |

implementation, making the sorting no longer the bottleneck for resource usage. This advantage is lessened when the settings are extreme. However, even when $S = 9$ and histogram size = 6, in which case the whole system can require too many logic cells to be implemented within our resources budget, the combinatorial one still uses fewer logic cells than the sequential version.

The remaining two modules, i.e. the Binner and Encoders, use few resources. These are normally below 100 LUTs+FFs each.

To guide the configuration selection, using only one encoder without sorting would be preferred because it can get rid of the histogram, sorter, mapper and encoder selector. If one has a histogram, increasing $S$ would be preferred over increasing histogram size because increased histogram size has a large effect on both the selector and histogram counter, which are the two resource-dominating modules. These findings are purely from the resource perspective, the selection should also be guided jointly on the resultant total power and BDP.

#### 7.2 Power consumption

Power consumption is another aspect of concern. We should guarantee that the added processing power does not exceed the reduced communication power. However, the estimated power indicates that the processing power consumption per channel is consistent among different configurations. As the histogram counter, sorter and encoder selector are only used during calibration, the Binner, Mapper, RAM and Encoder continuously consume energy. The binner and RAM tick at the processing clock/memory clock speed, but the input of the encoders only changes at $\frac{1}{BP}$ Hz, which is much lower than the clock speed. Therefore the power of the Binner and RAM dominate the FPGA dynamic power, which is around 0.96 $\mu$W per channel. The power of the encoder is negligible at 1 to 20 nW, the binner consumes about 0.46 $\mu$W per channel and RAM shares the remaining 0.5 $\mu$W per channel. For the remainder of this work, the combined compression/processing and communication power is referred to as the dynamic power. The board static power is 162 $\mu$W.

As all configurations use the binner and RAM, the processing power of different architectures is similar whether we encode the firing rate or not. Therefore the total power reduction we gain from the compression is proportional to the BR reduction.

Bases on the exploration of resources and power, we can conclude that it is the resources that constrain the algorithm complexity for the on-implant Huffman encoding.

### 8 Fixed Length vs. Variable Length Codewords and Bit-Flip Errors

An advantage of the multiplexed representation of MUA data is that the channel ID does not have to be explicitely encoded. This is because the channel ID is implicitely encoded in bit position. E.g.,

$$c = ceiling(k/m)$$

where $k \in [1 \leq k \leq n \times m]$ is the bit position and $c \in [1 \leq c \leq n]$ is the channel ID.

The principle technique of lossless compression is to transform fixed length codewords into variable length codewords. This is to take advantage of the potentially narrow and skewed nature of the data's histogram. However, fixed-length codewords have an advantage when it come to bit-flip errors during communication of multiplexed data. With fixed-length codes in multiplexed MUA, the channel ID is implicitly encoded in bit position, and this remains fixed over time. Therefore, any bit-flip error will only affect the communicated number of MUA events recorded on one channel, in the time period of question associated with the code block. No other channels will be affected.

With variable-length codewords, such as those produced by lossless compression techniques, all of the symbols after the codeword with the bit-flip error may be affected. Specifically, it may offset the relationship between symbols, encoders and channels. If different channels use different encoders, this can quickly make the entire sequence after the corrupted bit undecodable. For example in Table 1 (d), if the 3$^{rd}$ bit in the encoded multiplexed signal flips to a 1, then the sequence will be decoded as:

```
3, 2
```

instead of:

```
3, 0, 1
```

The consequences of the bit flip will propagate downstream. In this case, the 3rd symbol will be decoded as whatever comes next in the sequence, which should have corresponded to the 4th symbol. Given different channel-encoder pairings, the entire sequence post-error will likely become corrupted.

As such, if the bit-flip error rate is sufficiently high, some method may be required to reduce the consequences of bit flip errors. The first obvious candidate is to reduce the BP, where the BR is increased, but the temporal resolution of the data is increased. Therefore, if a data packet is corrupted, the user will not notice much difference as the next data packet will arrive shortly. The second clear candidate is noisy channel encoding.

#### 8.1 Noisy channel encoding

Noisy channel encoding involves adding parity bits to communicated data blocks. These parity bits have some relationship to the data. If a bit-flip error occurs, comparison of the codeblock and the parity bits can signal the existence of errors. The parity bits can even signal the locations of the errors, depending on the thoroughness of the noisy channel encoding and the number of parity bits.

If noisy channel encoding is used, the compression derived from variable-length codewords vs. fixed-length codewordsneeds to be sufficient to warrant the addition of parity bits to the code block.

In this work, noisy channel encoding was not explicitly applied as the rate of bit flip errors is unknown. However, calculating the required number of parity bits for each codeblock length is simple given the bit flip error rate and code block length.
